# Correlated Gene Modules Uncovered by Single-Cell Transcriptomics with High Detectability and Accuracy

**DOI:** 10.1101/2019.12.31.892190

**Authors:** Alec R. Chapman, David F. Lee, Wenting Cai, Wenping Ma, Xiang Li, Wenjie Sun, X. Sunney Xie

## Abstract

Single cell transcriptome sequencing has become extremely useful for cell typing. However, such differential expression data has shed little light on regulatory relationships among genes. Here, by examining pairwise correlations between mRNA levels of any two genes under steady-state conditions, we uncovered correlated gene modules (CGMs), clusters of intercorrelated genes that carry out certain biological functions together. We report a novel single-cell RNA-seq method called MALBAC-DT with higher detectability and accuracy, allowing determination of the covariance matrix of the expressed mRNAs for a homogenous cell population. We observed a prevalence of positive correlations between pairs of genes, with higher correlations corresponding to higher likelihoods of protein-protein interactions. Some CGMs, such as the p53 module in a cancer cell line, are cell type specific, while others, such as the protein synthesis CGM, are shared by different cell types. CGMs distinguished direct targets of p53 and exposed different modes of regulation of these genes in different cell types. Our covariance analyses of steady-state fluctuations provides a powerful way to advance our functional understanding of gene-to-gene interactions.

## Main Text

Single-cell RNA-seq (scRNA-seq) has greatly expanded our knowledge of gene expression. However, significant advances are still necessary to reach its full potential. Many methods have been developed for single cell amplification (*1-13*), but all suffer from various combinations of poor counting accuracy, low detection sensitivity, or low throughput. While these methods have been successful in cell typing (*5, 6, 14-17*), their ability to shed light on the roles of particular genes has been more limited. To further our understanding of how genes interact to produce complex cellular behaviors, a technique with high accuracy, sensitivity, and throughput is required. Such an understanding is critical to a wide range of biological problems, for example unraveling the networks of genes controlling cellular differentiation and identifying drug targets, to name only a couple.

To meet these unique technical demands, we designed a novel single-cell mRNA amplification method called Multiple Annealing and Looping Based Amplification Cycles for Digital Transcriptomics (MALBAC-DT) (Figure 1A). Our method improves upon several aspects of our prior work for amplifying DNA and RNA from single cells (*4, 18*). We improved transcript detection efficiency by optimizing reverse transcription to increase the amount of full length first-strand cDNA produced. First-strand cDNA then is amplified linearly by directly annealing MALBAC random primers along the cDNA strand, followed by exponential amplification by PCR.

**Fig. 1.**
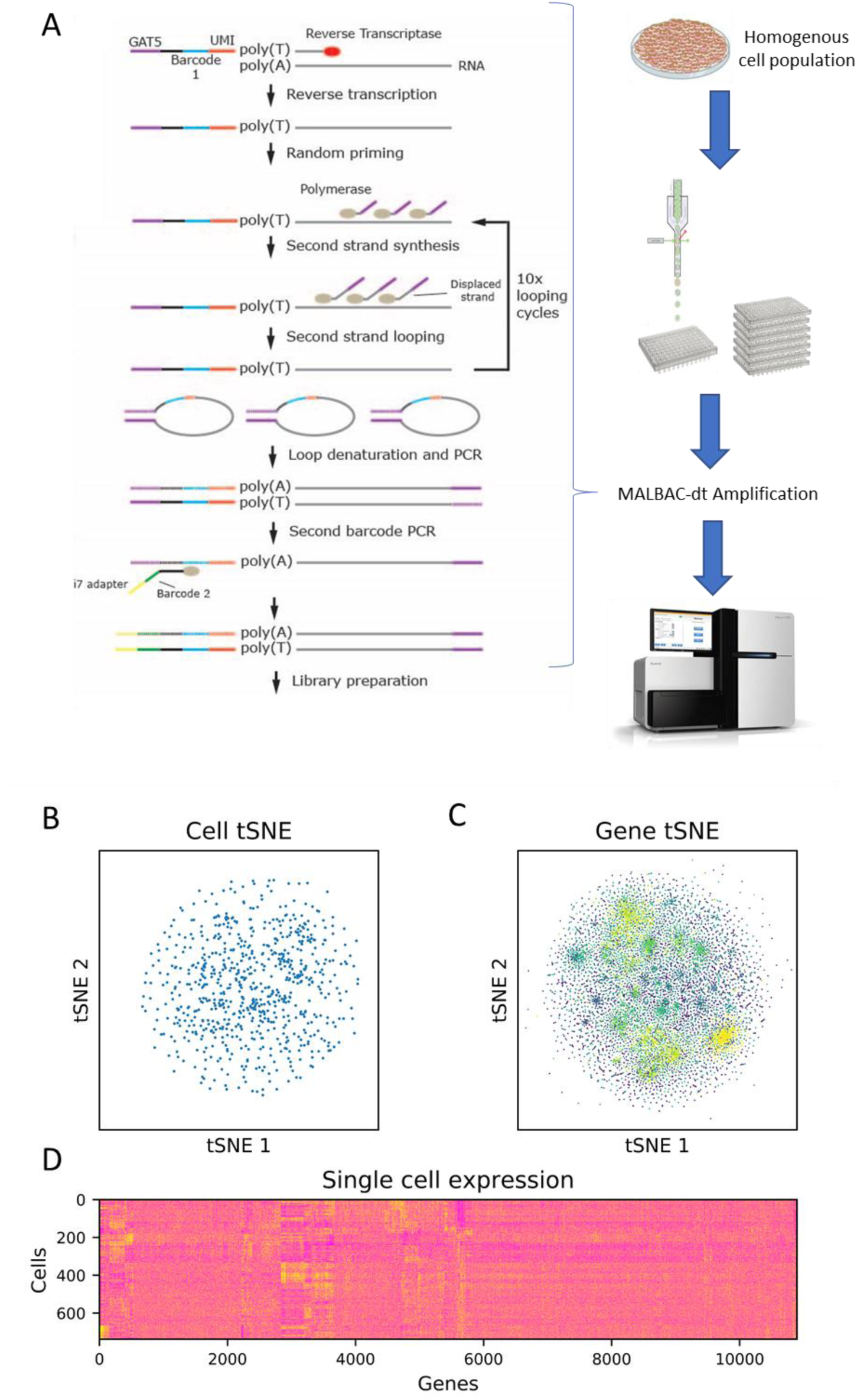
(A) MALBAC-dt protocol and experimental workflow. A homogenous cell population is trypsinized and sorted into individual wells of 96-well plates by flow cytometry. Reverse transcription is carried out using a poly-T primer containing a cell-specific barcode and unique molecular identifier (UMI). First strand cDNA is amplified by random primers using MALBAC thermocycling to ensure linear amplification followed by additional cycles of exponential amplification by PCR. After amplification, samples are pooled together for library preparation and sequencing. (B) Clustering of 738 U2OS cells by t-Stochastic Neighbor Embedding (tSNE). Cells were obtained from a homogenous culture and cDNA was amplified by MALBAC-dt. Consistent with a homogenous culture, no sub-clusters of cells are apparent. (C) Clustering of genes by tSNE. Genes were clustered based on their expression levels across the 738 U2OS cells. Many clusters are present, representing sets of genes that have similar expression patterns. (D) Hierarchical clustering of gene expression data from 738 U2OS cells. As with clustering by tSNE, several sets of genes with similar expression patterns are observed.

Because amplification by MALBAC-DT, in contrast to most single cell amplification methods, does not rely on template switching, we had greater flexibility to choose reverse transcriptases and optimize reaction conditions to maximize first strand cDNA production. Furthermore, with MALBAC-DT it is possible to successfully amplify transcripts that are only partially reverse transcribed, either due to their length or secondary structure. As a result, we detect ∼20% more genes from single-cell amounts of RNA compared to Smart-seq2 and obtain a lower percentage of reads corresponding to ERCC synthetic spike-ins which may be shorter or less complex than typical genes (Table S1).

To improve accuracy, we developed a novel unique molecular identifier (UMI) design that can correct previously unrecognized UMI artifacts that occur during amplification and sequencing (Supplementary methods). Although Smart-seq2 does not contain a UMI for absolute quantification of transcripts, we modified the protocol to incorporate the same UMI design to compare with MALBAC-DT and observed approximately twice and many transcripts detected when using MALBAC-DT (Table S2). Finally, our assay incorporates combinatorial cell barcoding to reduce the cost of preparing many single cells. Although we have opted to use UMIs and sequence only the 3’ ends of transcripts to improve quantification and reduce costs associated with library preparation and sequencing, we note that it is also possible to perform full length sequencing without UMIs by following standard library preparation protocols.

To demonstrate the ability of our method to generate unique insights into gene function, we amplified and sequenced 768 cells from the U2OS human osteosarcoma cell line, with 738 cells passing quality filters. As expected for a homogenous cell culture, clustering of cells based on gene expression using t-stochastic neighbor embedding (tSNE) (*19, 20*) did not reveal distinct subpopulations of cells (Figure 1B). However, clustering genes by tSNE did reveal distinct sets of genes that displayed similar patterns of expression across cells (Figure 1C). Clusters of genes that showed similar expression patterns across cells were also revealed by hierarchical clustering (Figure 1D).

To further investigate these clusters of genes, we computed the correlation coefficients for each pair of genes across all cells. Upon hierarchical clustering of the correlation matrix (Figure 2A), we observed 148 correlated gene modules (CGMs), or clusters of 10-200 genes that are highly correlated with each other. Many of these modules consist of genes pertaining to a specific biological function. These include general housekeeping functions—such as cell cycle control and cholesterol (Figure 2B) and protein synthesis (Figure 2C)—as well as functions pertaining specifically to this cell type such as bone growth and extracellular matrix remodeling (Figure 2D). A full list of CGMs and their associated functional enrichments is provided in Table S3.

**Fig. 2.**
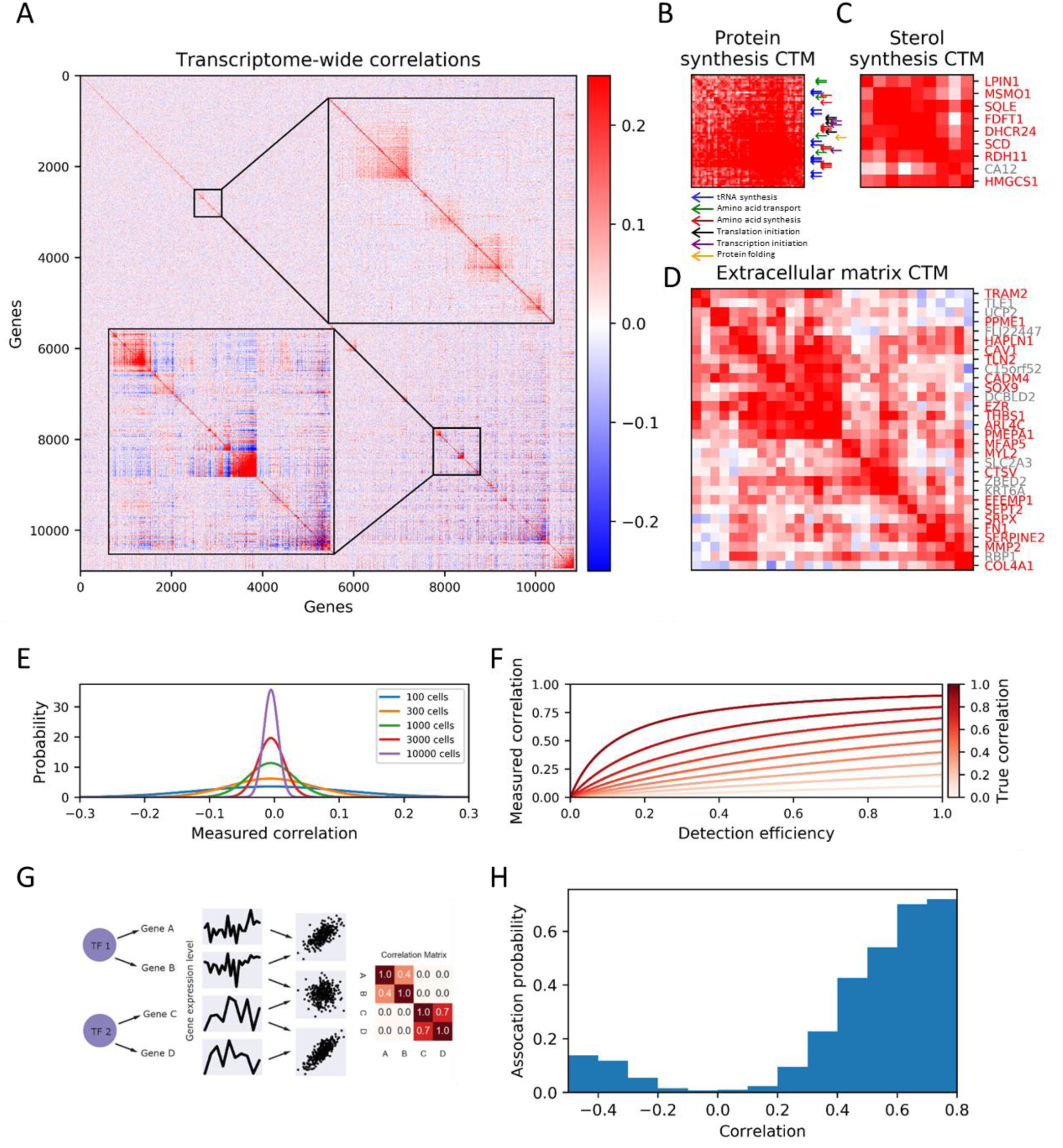
(A) Correlation matrix for ∼11,000 genes that were detected in at least 10% of the 738 U2OS cells. Genes are ordered by hierarchical clustering to reveal numerous modules of highly correlated and/or anti-correlated genes. Inset provides an enlarged view to highlight the detail present in many of these clusters. The genes in many of these Correlated Transcriptional Modules (CGMs) are enriched for particular biological function or contain binding sites for specific transcription factors. (B) A CGM related to protein synthesis, with genes responsible for specific processes indicated by arrows. (C and D) CGMs related to sterol synthesis (C) and extracellular matrix remodeling (D). Genes with known functional roles in these processes are labeled in red. (E) Measurement error associated with estimating correlation from a limited number of cells. The Plotted are the distributions of correlations that would be measured for a pair of uncorrelated genes if a given number of cells were sampled. (F) Impact of detection efficiency on correlation measurements. For a pair of genes with a given true correlation, the correlation that would be measured by sampling an unlimited number of cells is plotted as a function of the efficiency of detecting individual transcripts. (G) Genes with high correlations are more likely to be identified as related in protein-protein interaction databases. Gene pairs are binned based on their correlation coefficient. For each bin, the fraction of pairs identified by StringDb is plotted. (H) A schematic model depicting how gene regulatory interactions can result in correlations in a steady state population. Stochastic fluctuations in a transcription factor will result in fluctuations in its target genes, causing them to be correlated. Independently regulated genes, on the other hand, will exhibit no correlation.

It is evident that many CGMs have genes related to specific functions. For example the protein synthesis module (Figure 2B, Table S3 module 45), includes genes responsible for synthesizing tRNAs and amino acids, as well as the machinery required to initiate transcription and translation. The same is true for cholesterol synthesis (Figure 2C, Table S3 module 27). We note gene-to-gene correlation measurements have been widely used, but almost exclusively by means of the perturbative approach (*21-24*), i.e. evaluation of the correlations after introduction of a new experimental condition. This perturbative approach is bound to affect a large number of genes in the cell, usually resulting in large groups of correlated genes. Our method of evaluating correlations of steady state fluctuations of single cells reveals CGMs with a smaller number of intrinsically correlated genes.

While analysis and normalization of cell cycle dependencies is important for cell typing and differential expression analyses (*25*), the CGMs we observed are largely unaffected when expression levels are adjusted to control for cell cycle, with the exception of those CGMs that are directly related to cell cycle activity (Figure S1). Although genes representing different phases of the cell cycle become uncorrelated after normalization, we observe that genes within cell cycle related CGMs remain correlated, as would be expected due to correlations in the stochastic fluctuations that are not removed by normalization.

Detecting CGMs relies on precise measurements of correlations between genes. This requires large numbers of cells, accurate quantification of transcripts, and high detectability. To our knowledge, this is the first study that satisfies all of these criteria. Low cell numbers add sampling noise to the correlation coefficients (Figure 2E), while low detection sensitivity attenuates the correlations (Figure 2F). Simulations demonstrate that neither high cell numbers nor high detection sensitivity alone is sufficient (Figure S2-3), and that the CGMs detected were not the result of spurious correlations due to limited sample size (Figure S4). Previously published data (*8*) generated by 10x Genomics, which has high throughput but low sensitivity is unable to reveal CGMs (Figure S5), further confirming that large cell numbers cannot compensate for low sensitivity. A dataset more recently made available by 10x Genomics is able to detect a small number of modules (40 vs 178 CGMs identified by MALBAC-DT using the same cell line). Of these 40 modules, 53% (21/40) are found to be significantly correlated by MALBAC-DT (q < .05, Supplementary methods), while of the 178 CGMs identified by MALBAC-DT, only 9.6% (17/178) are able to be detected as significantly correlated in the 10x Genomics data.

Weighted Gene Correlation Network Analysis (WGCNA) has long been used to identify networks of genes using differential expression data across many cell types and conditions (*26, 27*). While this method of inference is logically distinct from our approach of using correlations within a uniform population, we found that the tools developed for WGCNA could also identify modules in our data. Adding further weight to the biological significance of the CGMs, the modules identified by WGCNA were highly similar to the CGMs we identified (Figure S6, Supplemental methods), although only 19 modules were identified by WGCNA. Modules such as tRNA aminoacylation, mitochondria, translation, and cell cycle are consistently found using both methods. However, modules like glycerolipid metabolism can only be identified with our analyses of the MALBAC-DT data.

Although it is widely understood that related genes will have similar differential expression levels across cell types or in response to perturbations, less consideration has been given to using steady state fluctuations to identify related genes. In general, the transcript levels of two genes under steady state conditions may be correlated if their transcription or degradation rates are correlated. This can arise from a number of biological mechanisms. Overall anabolic and catabolic activity of the cell will affect most genes indiscriminately but is largely removed by normalization. One gene-specific mechanism resulting in correlated transcription rates is coregulation either by a common transcription factor or multiple transcription factors which are themselves coregulated, possibly post-transcriptionally. Other possibilities include correlated changes in epigenetic states such as DNA methylation, histone modifications, or spatial position within the nucleus. Gene-specific mechanisms resulting in correlated degradation rates include common regulatory features in the untranslated regions (UTRs) or regulation by correlated miRNAs.

In light of these mechanisms that potentially result in gene correlations, we expect that genes that are involved in a common function, and hence are coregulated, would exhibit correlations such as those in Figure 2B-D. We examined whether a simple model of coregulation of gene expression is sufficient to produce the magnitude of correlations we observe in the CGMs. Due to the small number of DNA molecules present in a single cell, transcripts and proteins undergo stochastic fluctuations in expression level over time. When a regulatory protein controls the expression of multiple genes, it is natural to expect that fluctuations in the regulator will flow through to its targets, causing the targets to fluctuate in sync with one another. When their expression is measured across many cells, these genes would then be observed to be correlated (Figure 2H). Indeed, we find that conservative assumptions about the regulatory mechanisms underlying transcription and degradation are sufficient to produce correlations of the magnitude that we observe (Supplemental text, Figure S7). In our data, we observe significantly more positively correlated than negatively correlated pairs of genes, indicating that coregulation is a more widespread mechanism of regulation (Figure S8).

More generally, we observed that highly correlated genes were more likely to have been previously identified as being related in databases (*28*) of protein associations, including direct protein-protein interactions (PPI) and inferred functional relations (Figure 2G). Consistent with the observation of function-specific CGMs, this indicates that steady state correlations can indicate functional relationships between genes, and raises the possibility of identifying mechanistically related sets of genes from such measurements.

One CGM we chose for further investigation consists of targets of the key tumor suppressor protein p53 (Figure 3A). This CGM contains 197 genes, most of which have been identified by ChIP-seq studies to contain p53 binding sites. Moreover, all p53 targets identified by a previous ChIP-chip study (*29*) of this cell line are contained in this module.

**Fig. 3.**
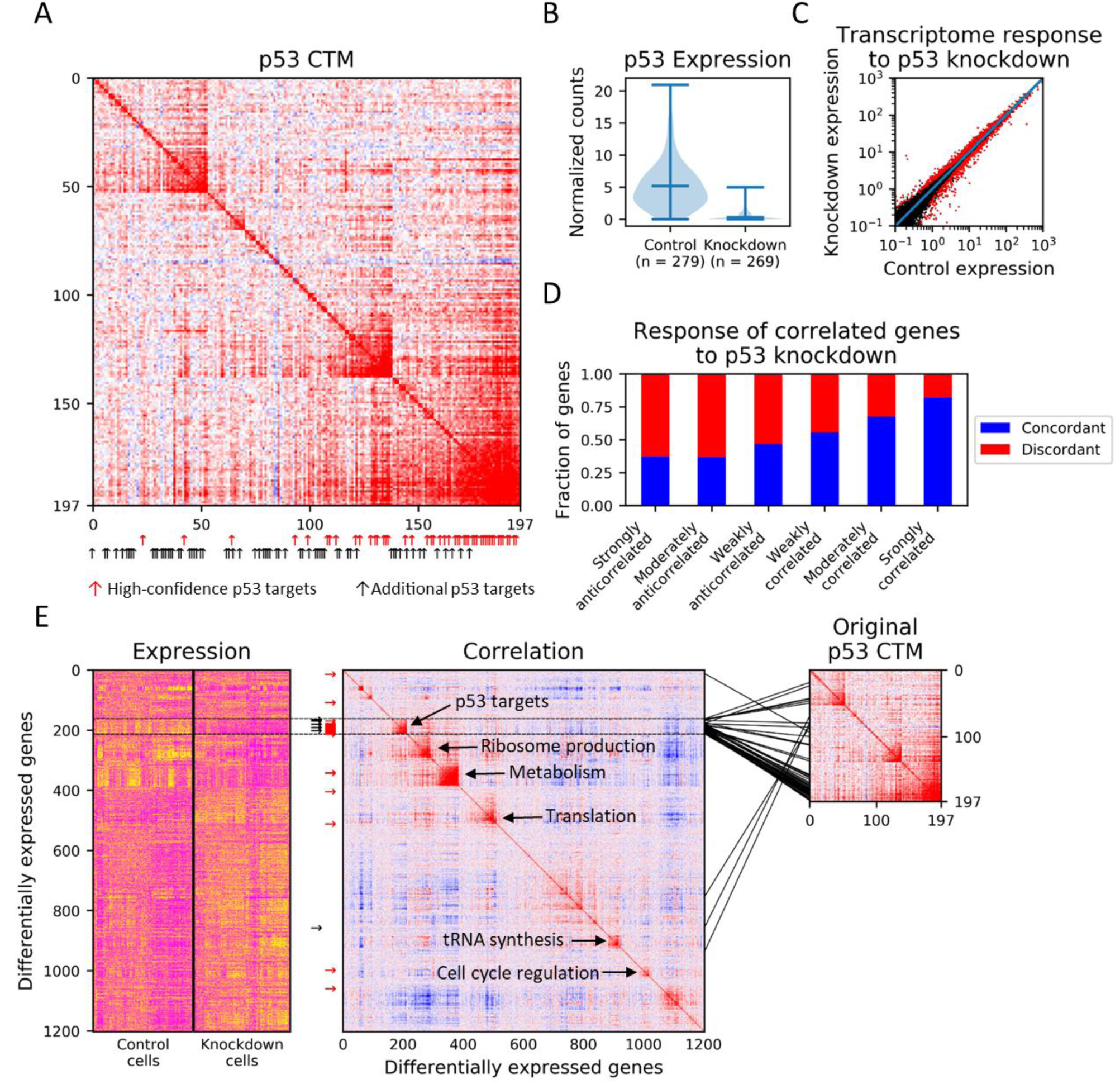
(A) CGM related to p53 activity. Genes with significant literature support (*31*) for being targets of p53 are indicated by red arrows, while genes with limited literature support are indicated by black arrows. (B) Distribution of p53 transcript levels in control and shRNA knockdown cells. (C) Mean expression levels of transcripts in p53 knockdown cells vs. control cells. Red points indicate genes which are significantly differentially expressed. (D) Correlation among genes in a homogenous cell population is predictive of their response to perturbation. Genes differentially expressed in response to p53 knockdown are categorized as strongly anticorrelated (less than −0.2), moderately anticorrelated (between −0.2 and −0.1), weakly anticorrelated (between −0.1 and 0), weakly correlated (between 0 and 0.1), moderately correlated (between 0.1 and 0.2), and strongly correlated (greater than 0.2). For each category, the fraction of gene pairs which are concordantly regulated (both upregulated or both downregulated) or discordantly regulated (one upregulated while the other downregulated) are shown. (E) Hierarchical clustering of genes differentially expressed upon p53 knockdown. Genes cluster into ∼10 CGMs. Genes within a CGM have the same directional response to p53 knockdown, consistent with their regulation as a functional unit. Direct p53 targets, indicated by red and black arrows as in (A), are predominantly found in a single CGM, as are the genes originally identified as a CGM related to p53 function. CGMs thus distinguish direct p53 targets from downstream pathways.

We performed an shRNA knockdown of p53 followed by MALBAC-DT in order to examine the effect of this module as well as the transcriptome as a whole. Mean p53 transcript levels decreased 15-fold in the knockdown cells compared to the control cells (Figure 3B), and 1337 genes were significantly up- or down-regulated as a result (Figure 3C).

If correlations between genes in our homogenous population of cells reflect coregulatory relationships, then we should expect that these genes would respond similarly to perturbations. Indeed, we find that genes that are strongly correlated in our original dataset tend to be either both upregulated or both downregulated in response to p53 perturbation, while anticorrelated genes tend to have opposite responses to the perturbation (Figure 3D).

Because correlations reflect regulatory relationships, they have the potential to identify novel genes that act in the same regulatory pathway. Indeed, several genes exhibited a high correlation with p53 expression and were perturbed by p53 knockdown, despite not previously being connected to p53 to our knowledge. As a concrete example, we observed that the deubiquitinase JOSD1 is strongly anticorrelated with p53 activity. Although JOSD1 had not been previously associated with p53, other deubiquitinases are known to modulate p53 activity either directly or through Mdm2. Moreover, a structurally related protein ATXN3 was recently shown to stabilize p53 via deubiquitination (*30*). We therefore hypothesized that JOSD1 might play a role in the p53 pathway. Indeed, JOSD1 was observed to be upregulated upon p53 inactivation, consistent with their anticorrelation in the unperturbed system, and indicating a possible negative feedback loop in which p53 inhibits JOSD1 transcription, while the Josd1 protein stabilizes p53.

The large number of genes that are differentially expressed in response to a perturbation often hinders meaningful analysis of such data, and providing meaningful classifications of these genes for further experiments remains an open challenge. CGMs potentially offer a way to organize these differentially expressed genes into fine-grained modular units. To this end, we looked at the correlations in our original dataset among the 1337 genes which were differentially expressed in response to p53 knockdown (Figure 3E).

These differentially expressed genes clustered into ∼10 CGMs, several of which are associated with distinct pathways. One of these CGMs consists almost entirely of genes from the original p53 CGM. Moreover, nearly all of the differentially expressed genes with p53 binding sites are contained in this CGM, indicating that by analyzing correlations we are able to distinguish the direct targets of p53 knockdown from its downstream effects. Strikingly, correlations in our steady-state dataset were able to predict the genes that would be perturbed by p53 knockdown more accurately than ChIP-seq studies. Whereas 49% of genes in the p53 module were downregulated upon knockdown of p53, this was only the case for 33% of genes from a consensus of ChIP-seq studies (*31*), and 9% of genes identified by a ChIP-chip study of the same cell line (*29*). Additionally, genes within CGMs are consistently upregulated or consistently downregulated upon p53 knockdown, in agreement with our model in which correlations among genes arise from coregulation.

Finally, we asked how CGMs compare across cell types. We amplified and sequenced 748 single cells from the HEK293T human embryonic kidney cell line. As expected for dramatically different cell types, several thousand genes were differentially expressed between these two cell lines (Figure 4A-B).

**Fig 4.**
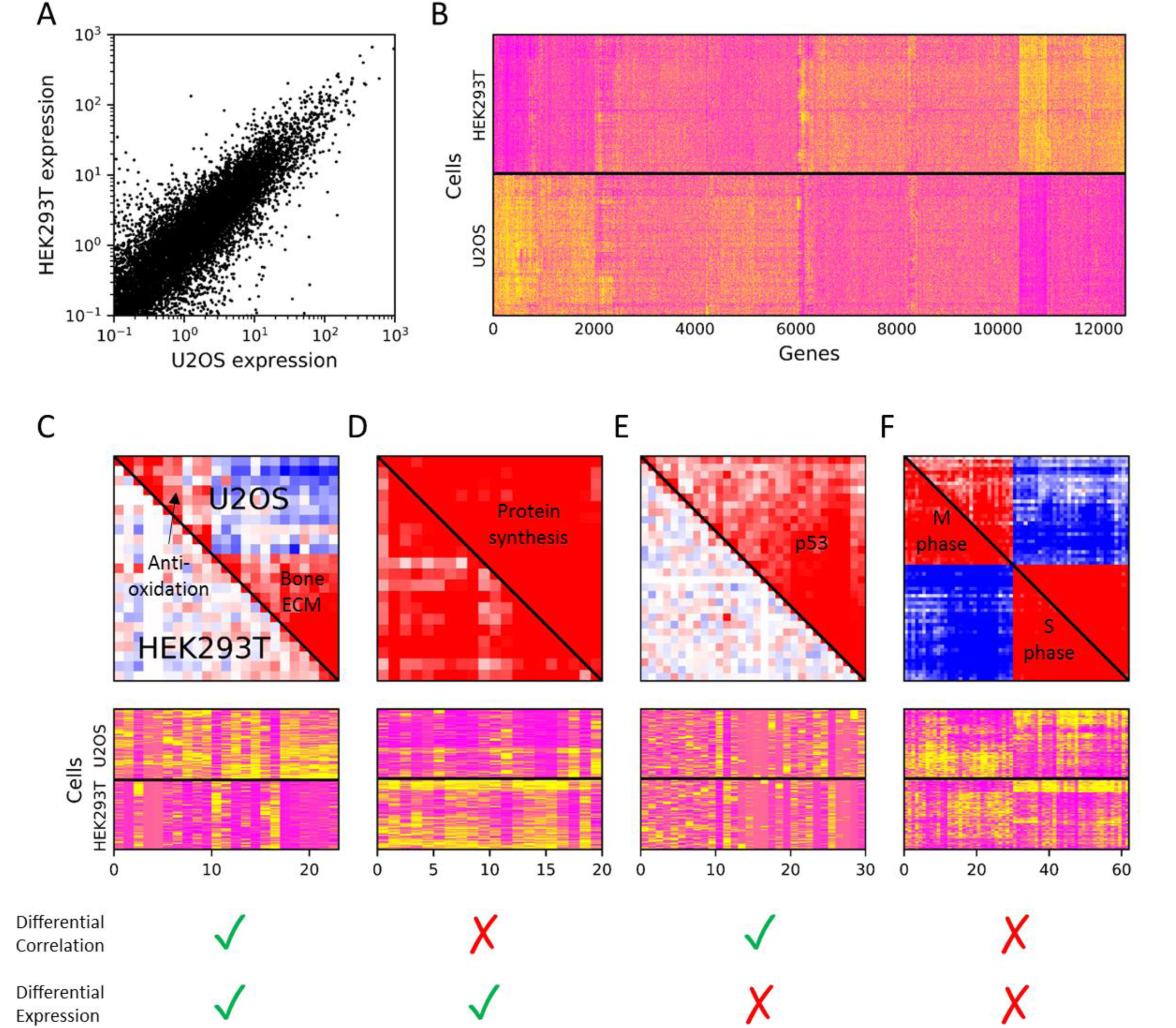
Shared and cell-type specific CGMs between the U-2 OS and HEK293T cell lines. (A) Mean expression levels of genes in HEK293T vs U2OS. (B) Hierarchical clustering of expression across HEK293T and U2OS cells reveals ∼6000 up- and down-regulated genes. (C-F) CGMs organize genes into biologically relevant pathways in a manner that is distinct from and not apparent by differential clustering. In some cases (C) differentially expressed genes correspond to differentially correlated modules. However, CGMs provide a further separation of differentially expressed genes into distinct functional units. In other cases (D), a set of differentially expressed genes can be resolved into a common CGM between multiple cell types. Moreover, differential correlation between cell types can occur without differential expression (E), possibly indicating multiple modes of regulation of the component genes. Finally, correlations can be consistently observed across cell types and organized into distinct CGMs in the absence of differential expression (F).

While existing methods are unable to provide meaningful classification of these genes, we find that CGMs provide a natural way to understand the differences between these two cell types. Of the 148 CGMs identified in U2OS, 22 CGMs were also observed to be significantly correlated in HEK293T cells, representing housekeeping machinery shared across vastly different cell types. For many CGMs, the genes are up- or down-regulated as a group (Figure 4C-D). In some of these cases, the CGM is absent in one cell type (Figure 4C), indicating that the module has been switched off. In other cases (Figure 4D), the CGM is present in both cell types, indicating that the genes remain coregulated but at different expression levels.

CGMs can also identify changes in regulatory architecture across cell types even in the absence of differential expression. Although HEK293T is known to lack p53 activity (*32, 33*), target genes of p53 are not consistently down-regulated in HEK293T compared to U2OS (Figure 4E), possibly indicating their roles in other pathways. Despite the similar expression levels of the component genes, the p53 CGM is absent in HEK293T.

Although in this work we have presented proof of principle analyses on cell lines using CGMs, our results shed light on critical biological processes relevant to many cell types. We expect the analysis of CGMs in a diverse set of cell types and tissues using methods with high sensitivity will produce a wealth of insights not obtainable using differential expression analyses alone.

## Acknowledgments

We thank L. Tan for helpful discussions of analysis methods and S. Mulepati for assistance with knockdown experiments.

## Author contributions

A.R.C., D.L., and X.S.X. designed the experiments. A.R.C., D.L., and W.M. performed the experiments. A.R.C. and W.C. analyzed the data. A.R.C., D.L., and X.S.X. wrote the manuscript.

## Data and materials availability

All sequencing data will be deposited in the NCBI SRA prior to publication.

## Supplementary Materials

Materials and Methods

Figures S1-S8

Tables S1-S3

## Supplementary Materials

### Cell culture and handling

U2OS and HEK293T cell lines were obtained from ATCC and cultured at 37°C in RPMI-1640 medium with 10% Fetal Bovine Serum and 1% Penicillin-Streptomycin. To form single cell suspensions for flow sorting, culture medium was removed, cultures were rinsed with Dulbecco’s phosphate-buffered saline (D-PBS), and incubated with 1mL of 0.25% trypsin for 5 minutes. Detached cells in D-PBS were pelleted by centrifugation at 300g for 5 minutes and resuspended in D-PBS. Single cell suspensions were kept on ice until flow sorting.

shRNA knockdowns were performed by incubating ∼30% confluent U2OS cells with 1ug of either 11653 C3 plasmid for TP53 knockdown cells or with TransIT-LT1 plasmid for control cells. Cells were incubated for 48 hours followed by flow sorting to isolate single cells in lysis buffer.

### MALBAC-DT protocol

Cells were flow sorted into 3uL of lysis buffer consisting of 1uL H2O, 0.6uL 5x SSIV buffer, 0.15uL 10% ICA-630, 0.8uL 5M betaine, 0.05uL SUPERase In, 0.2uL 50uM RT-An primer, and 0.2uL 10mM dNTP mix. Plates are stored at −80°C until ready for amplification. Plates are kept on ice while pipetting and vortexed and briefly centrifuged after all pipetting steps.

To perform reverse transcription, plates are incubated at 72°C for 3 minutes, then 1uL of RT mix is added consisting of 0.264uL H2O, 0.16uL 5x SSIV buffer, 0.2uL 100mM DTT, 0.152uL SUPERase In, 0.024uL 1M MgSO4, and 0.2uL SuperScript IV. Plates are incubated for 10 minutes at 55°C.

Next, excess reverse transcription primers are degraded by exonuclease digestion. 1uL of exonuclease mix is added consisting of 0.1uL ExoI buffer, 0.1uL H2O, 0.6uL ExoI, and 0.2uL 50uM RT-Bn primer. Plates are incubated for 30 minutes at 37°C and then 20 minutes at 80°C.

Amplification is performed by adding 24uL of amplification mix consisting of 18.64uL H2O, 3uL ThermoPol buffer, 0.4uL 10mM dNTP mix, 0.16uL 100mM MgSO4, 0.4uL 50uM GAT-7N, 0.4uL 50uM GAT-COM, and 1uL Deep Vent (exo-). The following thermocycle program is run:

| Step | Temperature | Time |
| --- | --- | --- |
| 1 | 95 | 5:00 |
| 2 | 4 | 0:50 |
| 3 | 10 | 0:50 |
| 4 | 20 | 0:50 |
| 5 | 30 | 0:50 |
| 6 | 40 | 0:45 |
| 7 | 50 | 0:45 |
| 8 | 65 | 4:00 |
| 9 | 95 | 0:20 |
| 10 | 58 | 0:20 |
| 11 | Goto 2 | 10x |
| 12 | 95 | 1:00 |
| 13 | 95 | 0:20 |
| 14 | 58 | 0:30 |
| 15 | 72 | 3:00 |
| 16 | Goto 13 | 17x |
| 17 | 72 | 5:00 |
| 18 | 4 | 0:00 |

Finally, amplification is completed by adding 0.4uL 50uM Tru2-Gn-RT primers and running an additional 5 cycles of PCR steps 12-15. Amplified plates are stored at −20°C until library preparation.

In early versions of the protocol, a total of 8 RT3-An primers were used per 96 well plate, with one primer corresponding to one row of the plate. During the final amplification step, 12 Tru2-Gn-RT primers were used, with one primer corresponding to one column of the plate. In later versions of the protocol, 96 RT3-An primers were used, with a distinct primer corresponding to each well. In the final step, a single Tru2-Gn-RT primer could then be used. While the former method requires lower upfront costs to synthesize primers, the later method simplifies preparing plates at larger scales and eliminates the possibility of cross-contamination of samples during amplification.

To prepare libraries for sequencing, 1uL from each of the wells are combined and purified using 0.8x Ampure beads. The Nextera library preparation kit is used to add Illumina adapters by tagmentation. During subsequent PCR steps, Ix-Tru2 primers are substituted for Nextera S5XX primers in order to select the 3’ ends of transcripts containing cell barcodes and UMIs.

### Sequence processing

Separate fastq files are generated for each cell based on the outer and inner barcode sequences. Barcodes not matching a cell exactly are discarded. Barcodes, adapter sequences, and UMIs are stripped from the reads, and reads are aligned to the human GRCh38.p7 reference using STAR 2.5.2. For each gene, a list of UMIs is obtained for all reads mapping to that gene, excluding regions masked by RepeatMasker. To remove extraneous UMIs resulting from amplification or sequencing errors, UMIs for a particular gene are represented as nodes in a graph, with connections between UMIs differing at no more than 7 bases. Connected components are identified, and the consensus sequence within each component is determined. Consensus sequences matching the (HBDV)_5_ RT-An pattern and differing from the (VDBH)_5_ RT-Bn pattern at at least three bases are retained. To avoid potential cross-talk between wells, UMIs observed for the same gene in multiple cells are discarded.

After obtaining UMI counts for all genes and cells, cells for which more than 1% of transcripts are from ERCC spike-ins or contain fewer than 1000 total transcripts are discarded, as are genes which are observed in fewer than 10% of cells. Counts are normalized relative to the total number of transcripts in each cell prior to computing the correlation matrix.

Hierarchical clustering is performed using the SciPy function scipy.cluster.hierarchy.linkage using method “average,” and with a distance metric of 1 − *abs*(*ρ_ij_*), where *ρ_ij_* is the correlation between genes *i* and *j*. To test the robustness of this clustering algorithm, we randomized the umi counts across all cells for each particular gene and recomputed the correlations between gene pairs, resulting in a distribution of correlation coefficients that would be expected due to limited sample size alone. On top of this background of uncorrelated genes, we set a group of genes to have a stronger correlation and examined whether these genes could then be identified by clustering. We found that for the magnitudes of correlations typically observed in our data, groups of correlated genes could be reliably recovered (Figure S9).

### Cell cycle correction

Pseudo-time inference and cell-cycle correction

Pseudo-time was inferred for each cell by assuming that the expression of cell-cycle genes followed a sinusoidal function along the time trajectory. The actual expression of each cell-cycle gene was further modeled as follows, a normal distribution centered around the level predicted by sinusoidal function, with variance aggregated from both stochastic expression variance and technical noise.

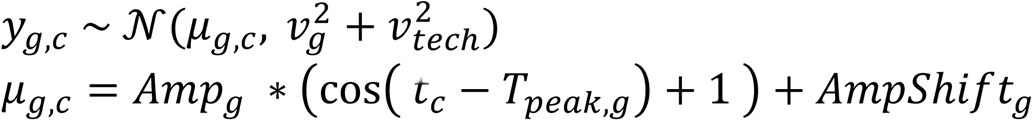

*y_g,c_*: actual expression of gene g for cell c.

*μ_g,c_*: expected expression of g for c from sinusoidal function.

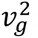: gene specific variance from stochastic expression for g.

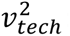: common technical noise.

*Amp_g_, AmpShift_g_*: amplitude of the sinusoidal function for g.

*T_peak,g_*: The peak time of g, in the time scale of percentage into the cell-cycle. Retrieved from Cyclebase.org (*34*).

*t_c_*: The pseudo-time of cell c.

The transcriptome was fitted against the described model, with a pseudo-time optimized for each cell to maximize the overall likelihood estimation. The MLE process was done using PyTorch.

In order to correct the covariance matrix for cell-cycle effect, cells were then ordered by the assigned pseudo-time, and the expression of each gene was corrected by subtracting the mean of the surrounding rolling window.

### Correlations between coregulated genes

We consider the case of two genes that are regulated by the same transcription factor (Supplementary Figure 1A) with dynamics described by:

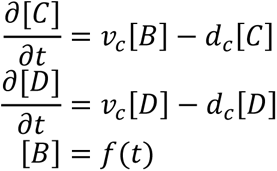

For simplicity we have taken the transcription rate *μ* and degradation rate *ν* to be the same for transcripts C and D. The transcription factor B is allowed to fluctuate in time arbitrarily.

The lifetimes of mRNAs are often short relative to those of proteins. In this case, as B fluctuates, transcripts C and D rapidly adjust and fluctuate independently from one another around the steady state concentration [*C*]*_ss_* = [*D*]*_ss_* = [*B*] *ν_c_*/*d_c_*, and these fluctuations will follow a Poisson distribution.

Under these assumptions, the covariance between [*C*] and [*D*] is:

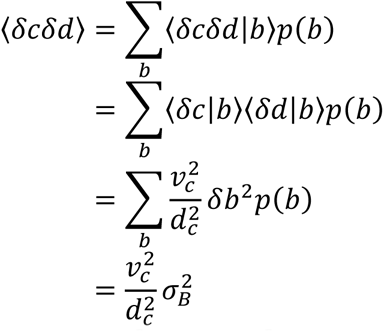

Following a similar procedure to obtain <*δc*^2^> and <*δd*^2^>, we obtain the correlation coefficient

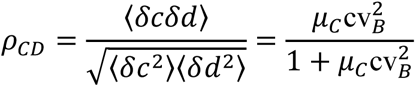

where *μ_C_* is the mean of C, and cv*_B_* is the coefficient of variation of B.

### Analysis of 10x Genomics datasets

Datasets were downloaded from the website of 10x Genomics. The datasets from Zheng et al (8) consisting of ∼2800 HETK293T cells and the newer dataset consisting of a mixture of ∼10,000 HEK293T and mouse NIH3T3 cells were used. For the later dataset, only HEK293T cells were used for further analysis. BAM files were downloaded and filtered according to the same mappability criteria used for MALBAC-DT datasets. To determine whether a CGM detected with one method was significantly correlated in the other, genes within a given module were randomly substituted for genes with similar expression levels (among the 50 genes with nearest mean expression), and the average of the absolute value of the correlations in this randomized module were calculated. This randomization was repeated 10,000 times and a Bonferoni-corrected p-value was obtained by comparing the average correlation in the true module to the distribution of average correlations in the randomized modules.

### Comparison to WGCNA

Hierarchical clustering of the correlation matrix is performed using the “hclust” function of R software, with the “average” method and a distance metric of 1-abs(ρ_ij), where ρ_ij is the correlation between genes i and j. Sub-clusters are obtained by cutting the dendrogram using the “cutree” function with parameter h=0.9. Sub-clusters containing more than 10 genes are identified as CGMs. We also compare our module detection method with a widely used gene co-expression analysis package, WGCNA1,2. Modules are identified by the “blockwiseModules” function, with “power = 1, TOMType = “unsigned”, minModuleSize = 10” and the other default parameters. Gene set enrichment analysis is performed using the R package “enrichR” with p-value threshold of 1e-5 to associate gene sets in modules with “KEGG_2019_Human”, “GO_Biological_Process_2018”, “GO_Cellular_Component_2018”, “GO_Molecular_Function_2018”, and “Reactome_2016” databases. For the U2OS cell line, only 19 modules are identified by WGCNA, and nearly 9,000 genes do not form any module. We use the Dice coefficient as a measure of similarity between modules detected by both methods and found that CGMs and WGCNA modules are highly similar.

## Supplementary Figures

**Figure S1.**
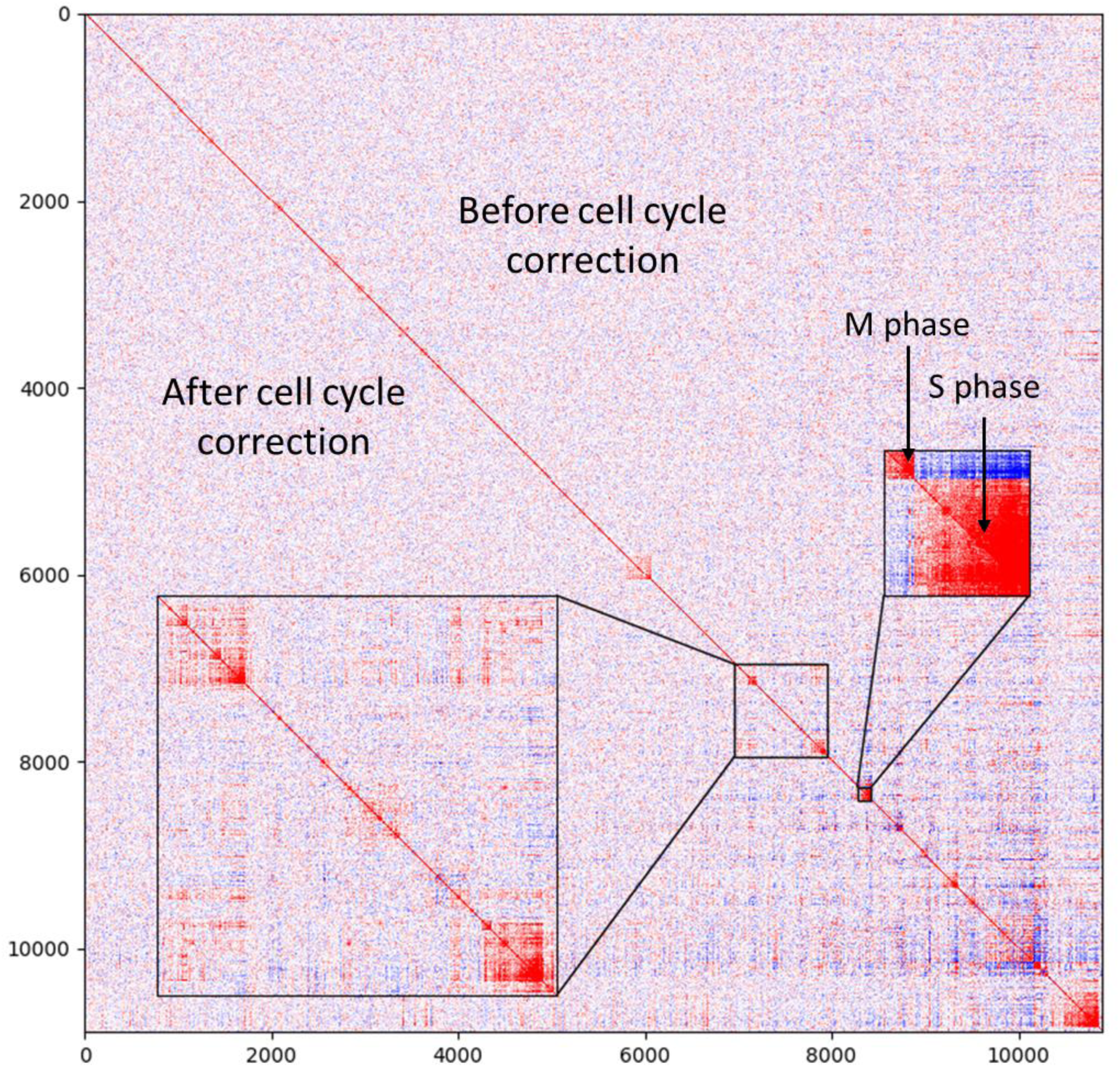
Effect of cell cycle on CGMs. Correlations are shown before (upper right) and after (lower left) adjusting expression data for cell cycle differences across cells. Nearly all CGMs are unaffected by the correction, with individual pairs of genes exhibiting similar correlations before and after adjustment. Genes specifically related to the cell cycle are the exception. Related cell cycle genes generally exhibit weaker correlation after adjustment, and the negative correlations between M and S phase genes are largely eliminated.

**Figure S2.**
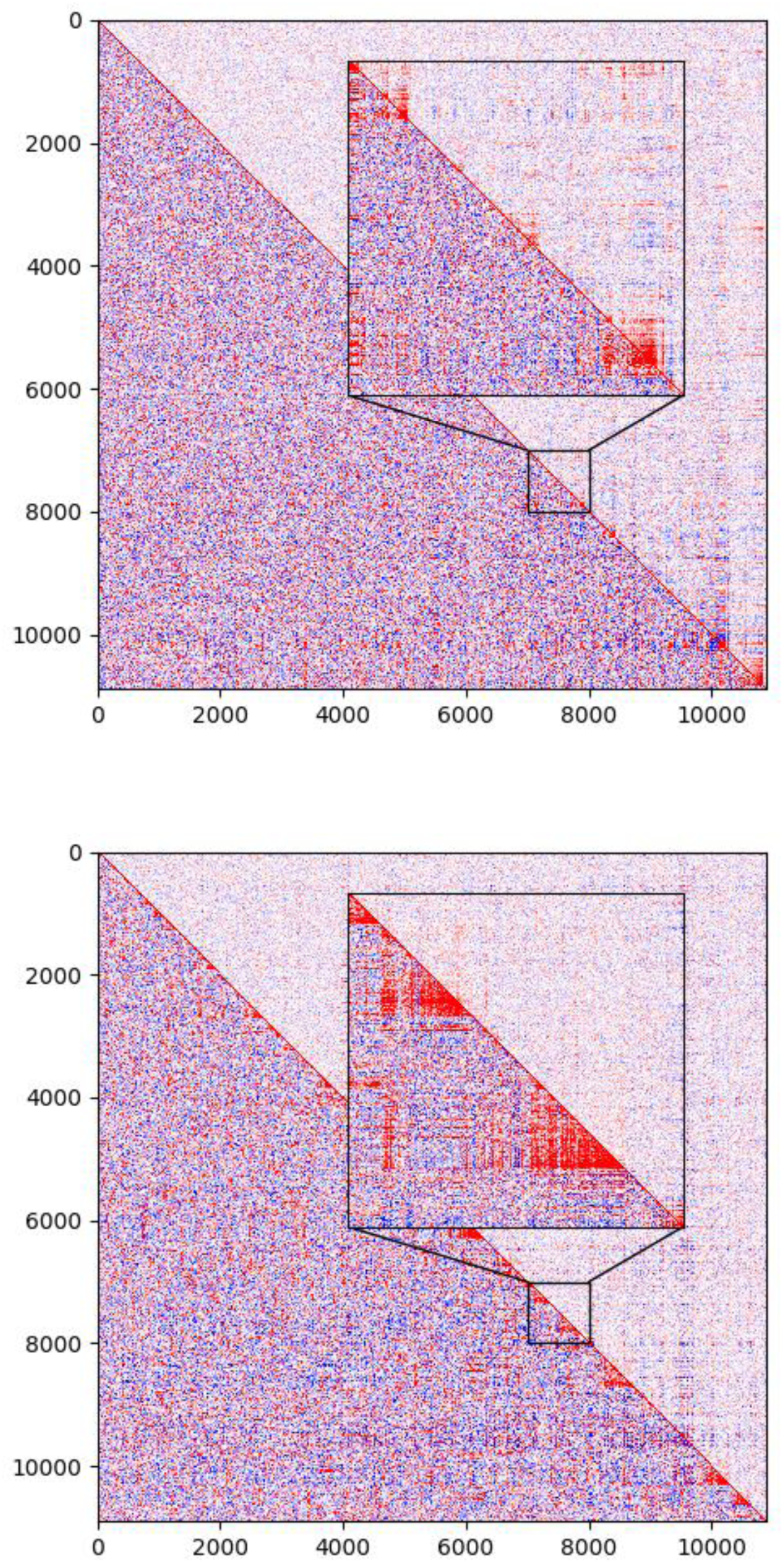
Large numbers of cells are needed to observe CGMs. Correlations in the full U2OS dataset (upper right) are compared with correlations calculated using a random subset of 100 cells (lower left). The 100 cell subset is significantly noisier than the full dataset. In many cases, CGMs identified in the full dataset (A) are observed to also be correlated in the 100 cell subset. However, the larger noise in this dataset prevents CGMs from being identified using this reduced dataset alone. When genes are clustered according to their correlation in the 100 cell dataset (B), many spurious clusters are identified. The correlations within such clusters result solely from measurement error and are not observed in the larger dataset, which has lower measurement error due to the larger number of cells.

**Figure S3.**
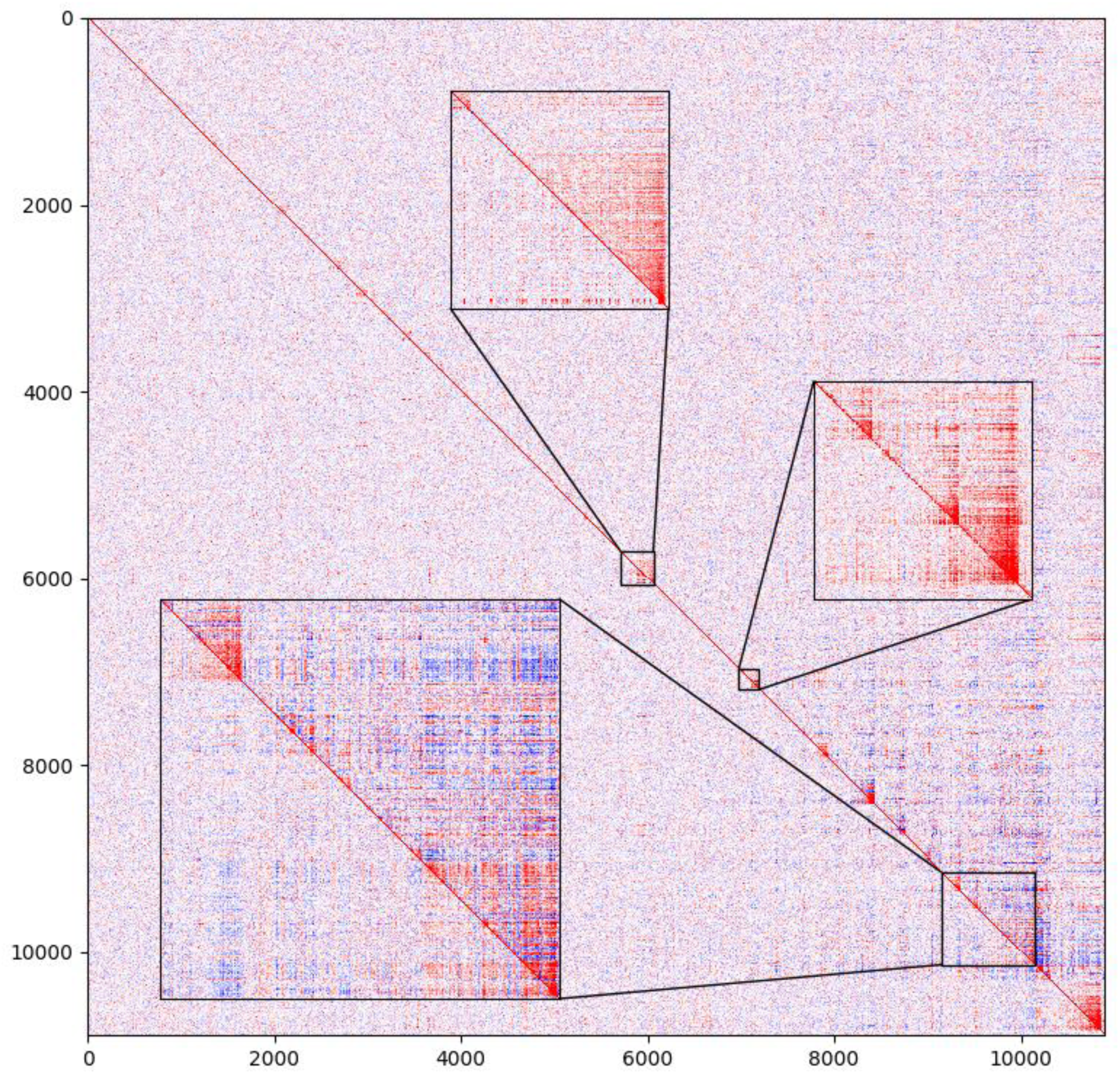
High detection efficiency is necessary to detect CGMs. Correlations between genes are shown for the full U2OS dataset (upper right), and a randomly downsampled dataset simulating a 67% lower detection efficiency (lower left). Many correlations are absent or severely attenuated at lower detection efficiency.

**Figure S4.**
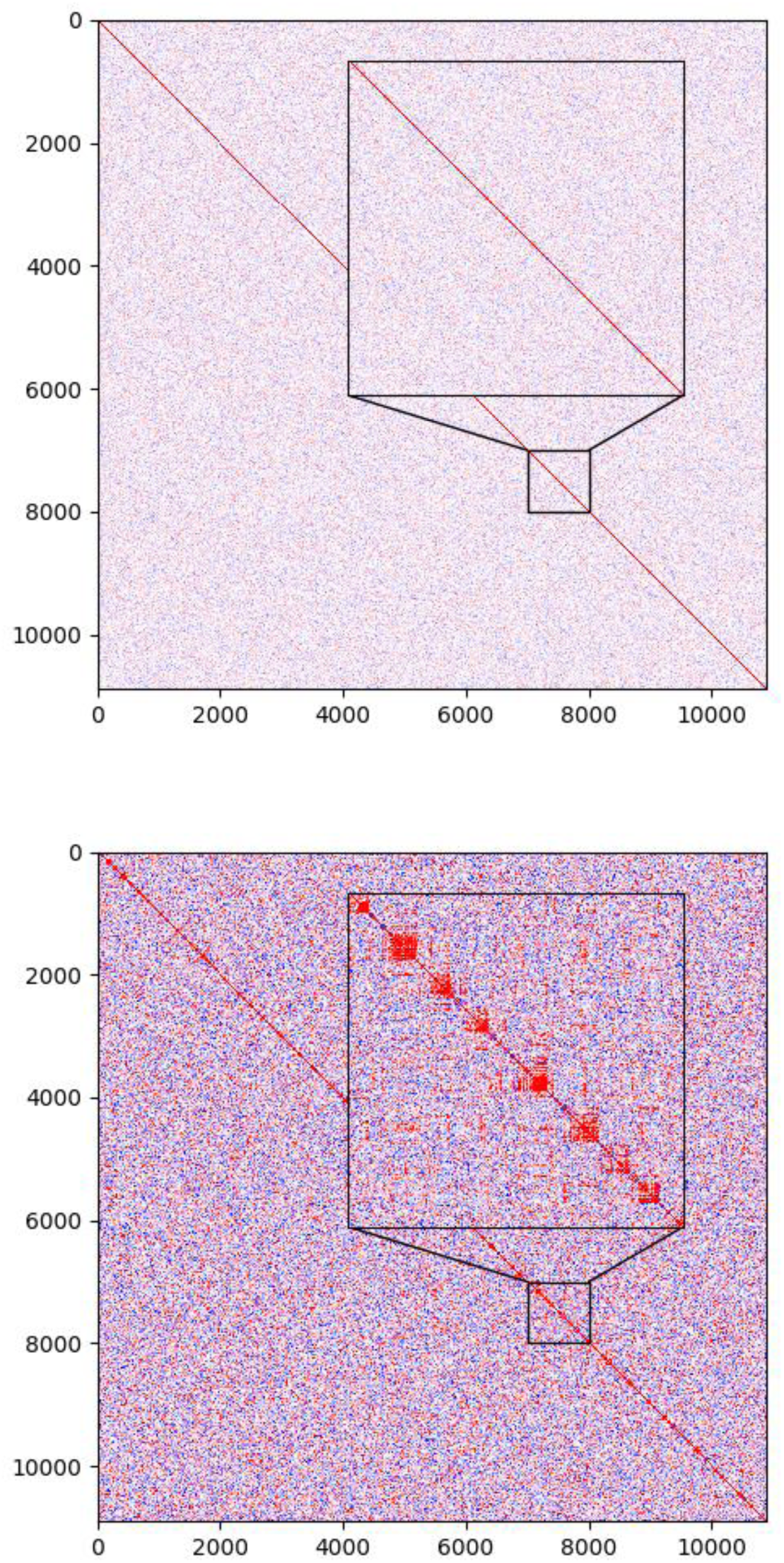
CGMs do not result from measurement error. To examine the effect of measurement error in the correlation coefficients on clustering, we randomly permuted the expression counts across all cells for each gene. As a result, there is no expected correlation between genes, and any observed correlation is due to sampling a finite number of cells. In our full U2OS dataset (A), no CGMs are observed after random permutations. Spurious clusters are observed when the permutation is applied to a random subset of 100 cells (B).

**Figure S5.**
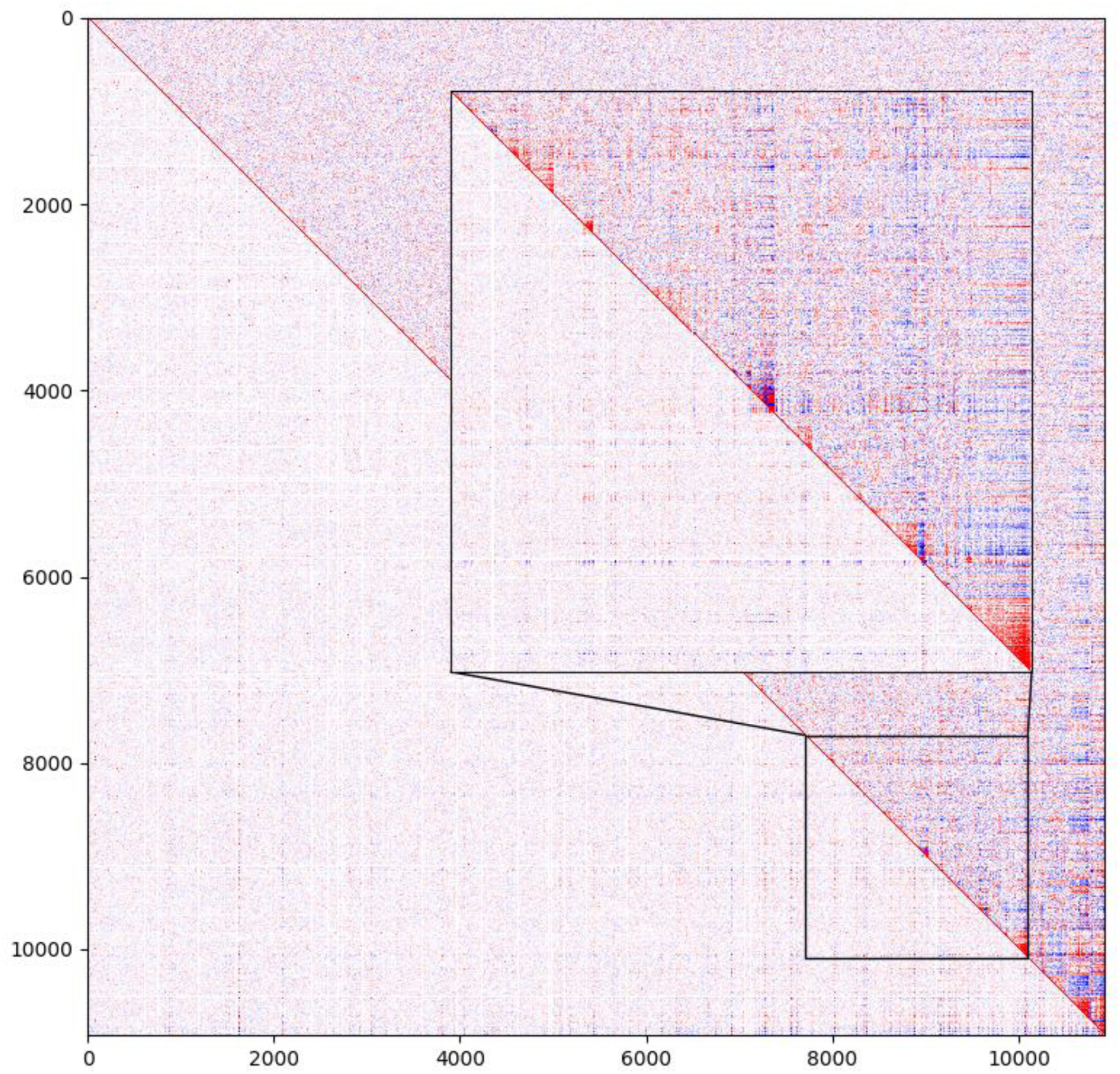
Methods with low sensitivity cannot detect CGMs. Correlations are displayed for data generated by MALBAC-DT (upper right) and 10x Genomics (lower right) for the HEK293T cell line. Although 2800 cells were sequenced with 10x compared to 748 cells with MALBAC-DT, correlations and CGMs are only apparent in the MALBAC-DT data.

**Figure S6.**
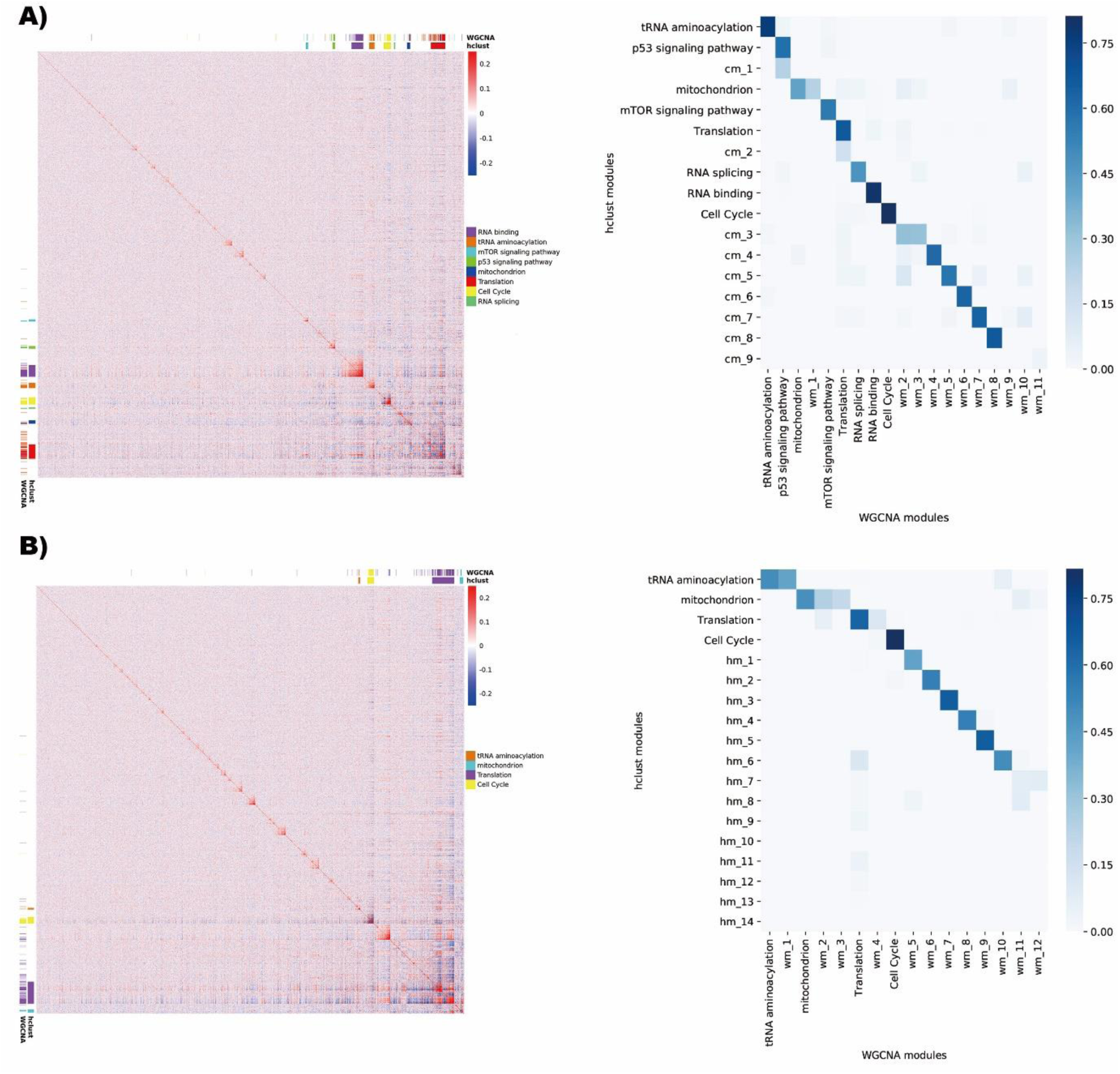
CGMs are consistent with the modules detected using WGCNA. A) Left panel: hierarchical clustering of gene-gene correlation across U2OS cells. Modules associated with specific functions are highlighted using color bars for both methods. Right panel: heatmap of Dice’s coefficient indicates the similarity between the CGMs and the WGCNA modules. Modules associated with specific functions are detected by both methods with shared gene sets over 40% of the average number of genes in the two modules. 84% of the modules detected by WGCNA can be found in CGMs with Dice’s coefficient > 0.4. B) Hierarchical clustering of gene-gene correlation across HEK293T cells. The four function-associated modules share gene sets over 40% of the average number of genes in the two modules. 87.5% of the modules detected by WGCNA can be found in CGMs with Dice’s coefficient > 0.4.

**Figure S7.**
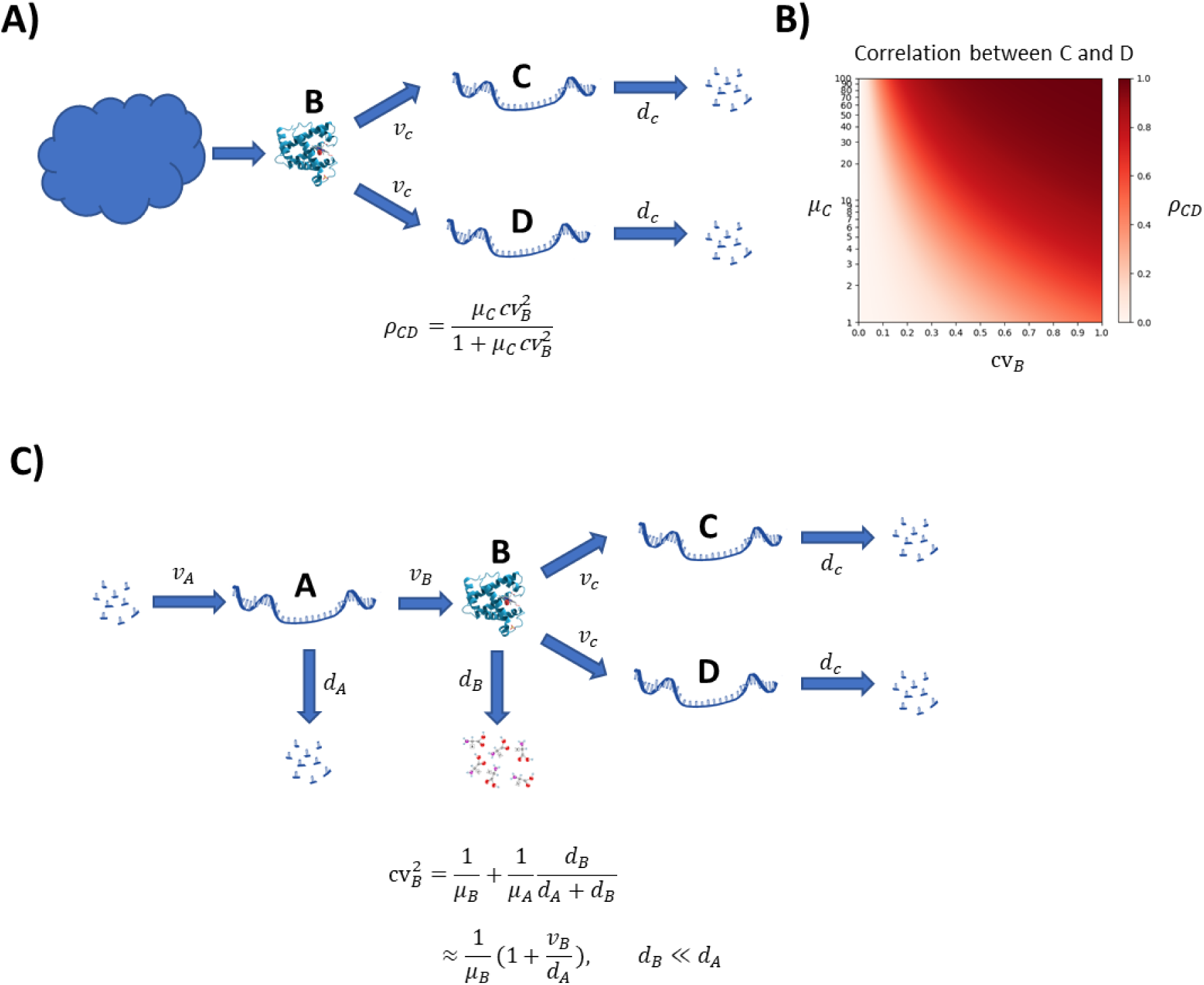
Model of correlations resulting from shared transcriptional regulation (see also supplementary note). A) A transcription factor B regulates the transcription of genes C and D, with the rate of transcription given by [B]*ν_C_*. Transcripts C and D are degraded with rate *d_C_*. The protein B fluctuates in time, driven by an arbitrarily complex mechanism regulating its production and degradation. Under the assumption that the mRNA lifetimes of C and D are short relative to the timescale of fluctuations of B, the correlation between the two genes is given by 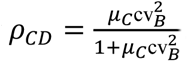. B) Heatmap showing the magnitude of correlation coefficients that are obtained for various values of the coefficient of variation of B (cv*_B_*) and mean expression level of C and D (*μ*_C_). C) Correlations under a particular model in which fluctuations in B are driven by a simple model of transcription and degradation at constant rates with no bursting. In this case, under the assumption that the protein B is long-lived relative to its transcript A, the coefficient of variation is given by 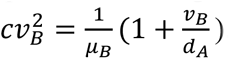.

**Figure S8.**
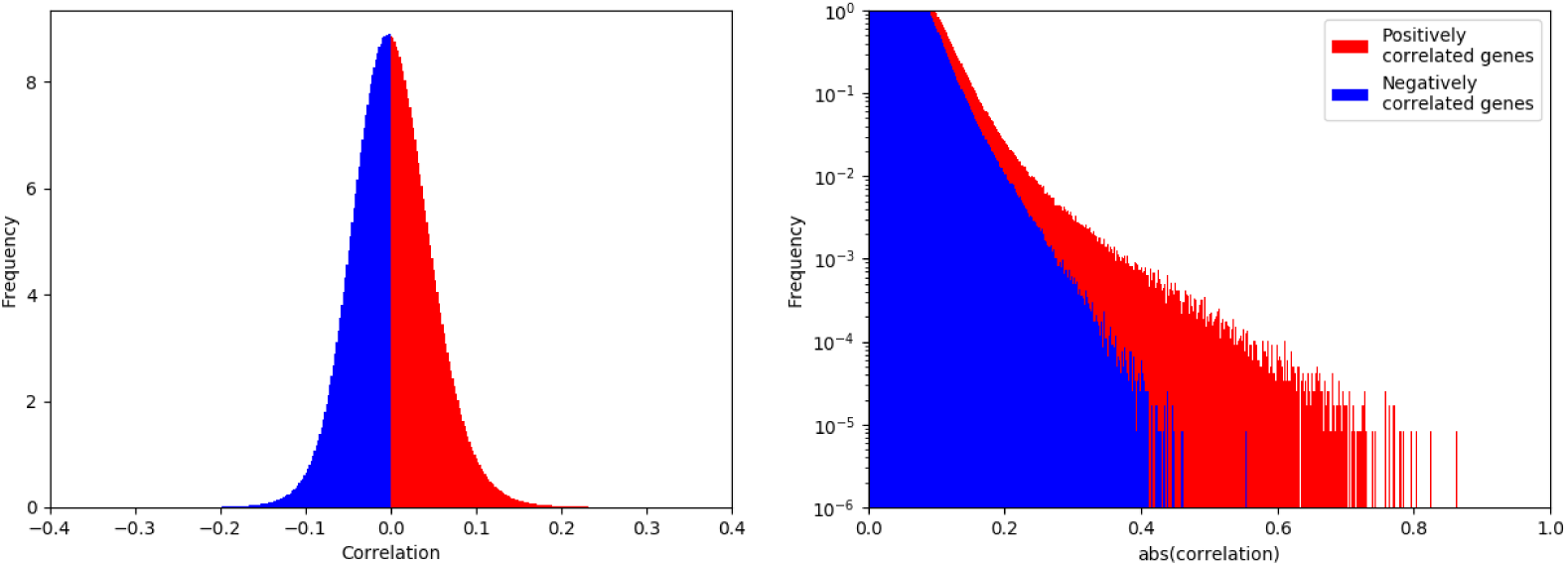
Distribution of correlation coefficients. A) Histogram of measured correlation coefficients for all pairs of genes. The portion of the distribution corresponding to genes with positive correlation is show in red, and the portion corresponding to negatively correlated genes is shown in blue. The distribution is roughly symmetrical with most gene pairs being uncorrelated. B) Zoomed in view of panel (A) highlighting the tails of the distribution. Negative correlations are displayed on the positive axis for better contrast with positive correlations, and the frequency is displayed on a log scale. Strongly correlated gene pairs are significantly more prevalent than strongly anticorrelated gene pairs.

**Figure S9.**
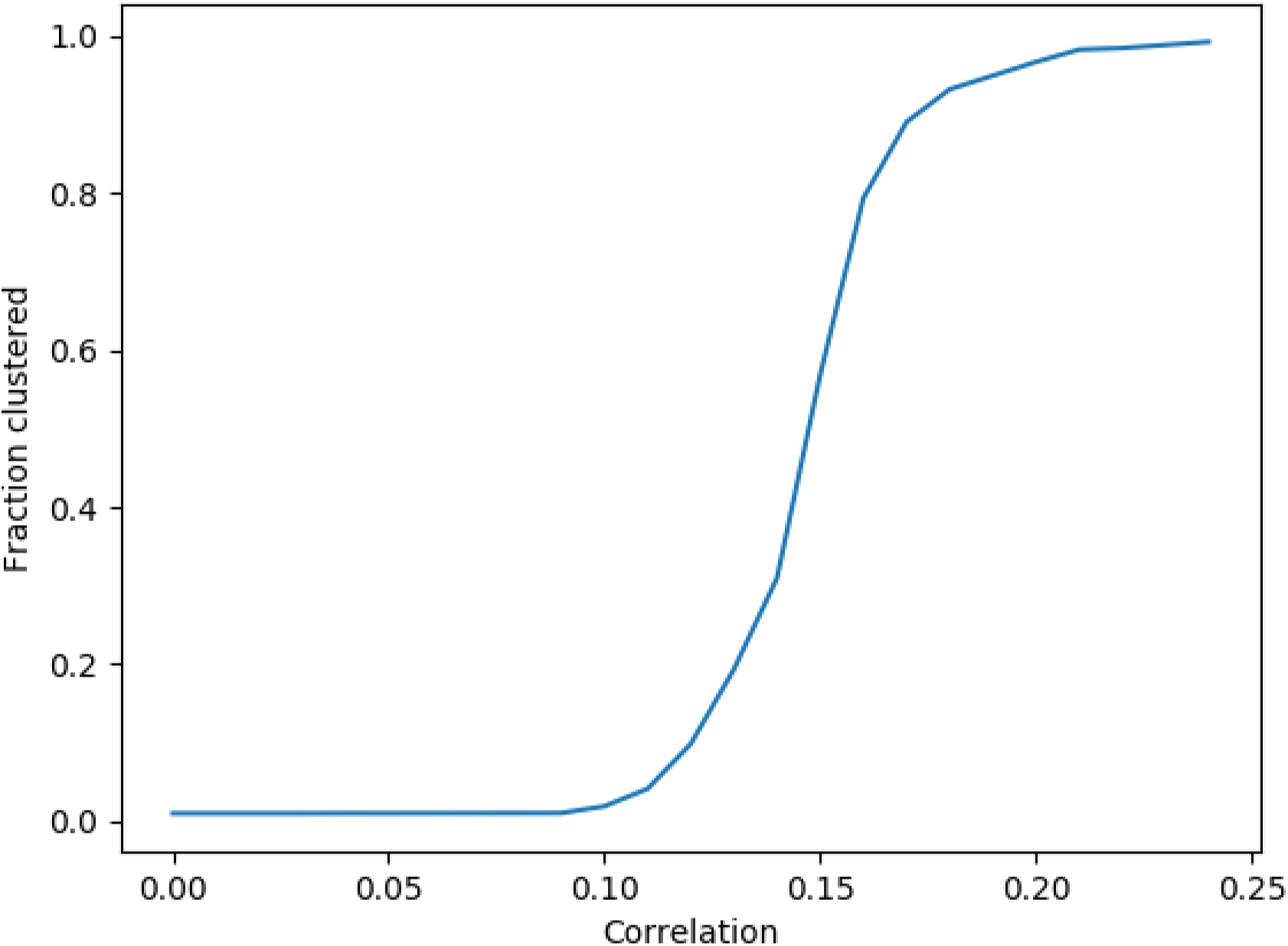
Clustering is robust at the magnitudes of correlation observed in our data. A set of 100 genes are set to have a fixed correlation against a background of correlations due entirely to sampling error. Clustering is performed, and the fraction of pairs of genes which are in the same cluster is recorded. The mean fraction is plotted across 10 simulations over different random backgrounds.

## Supplementary Tables

**Table S1.** Performance characteristics of MALBAC-DT compared to Smart-seq2. All samples have been downsampled to 250,000 mapped reads for comparison. In order to prevent inclusion of ambiguously mapped reads and inflated gene counts, reads have been stringently filtered to exclude regions flagged by RepeatMasker.

| Sample | Mapped reads | Exonic fraction | ERCC fraction | Genes detected |
| --- | --- | --- | --- | --- |
| MALBAC-DT Replicate 1 | 250000 | 89% | 0.2% | 4984 |
| MALBAC-DT Replicate 2 | 250000 | 88% | 0.2% | 5567 |
| Smart-seq2 Replicate 1 | 250000 | 92% | 2.6% | 4326 |
| Smart-seq2 Replicate 2 | 250000 | 93% | 2.8% | 4605 |

**Table S2.** Performance of MALBAC-DT compared to a modified Smart-seq2 protocol containing the same UMI design as for MALBAC-DT. All samples have been downsampled to 380,000 mapped reads for comparison, and the number of transcripts is presented after correcting for amplification and sequencing artifacts.

| Sample | Mapped reads | Transcripts detected |
| --- | --- | --- |
| MALBAC-DT Replicate 1 | 380,000 | 48,389 |
| MALBAC-DT Replicate 2 | 380,000 | 47,736 |
| MALBAC-DT Replicate 3 | 380,000 | 47,711 |
| MALBAC-DT Replicate 4 | 380,000 | 49,830 |
| Smart-seq2 with UMIs, Replicate 1 | 380,000 | 24,664 |
| Smart-seq2 with UMIs, Replicate 2 | 380,000 | 20,946 |

**Table S3.**
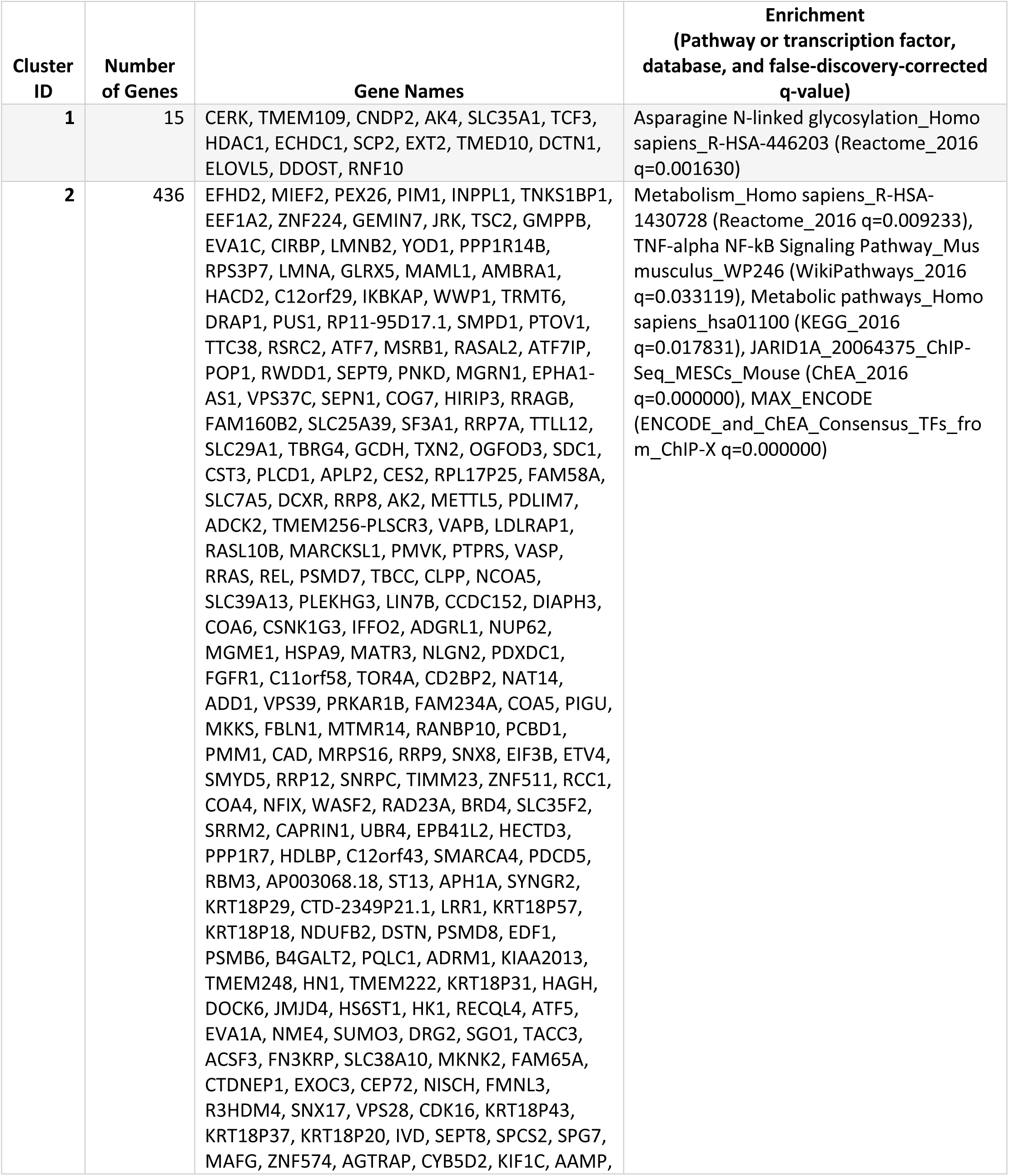
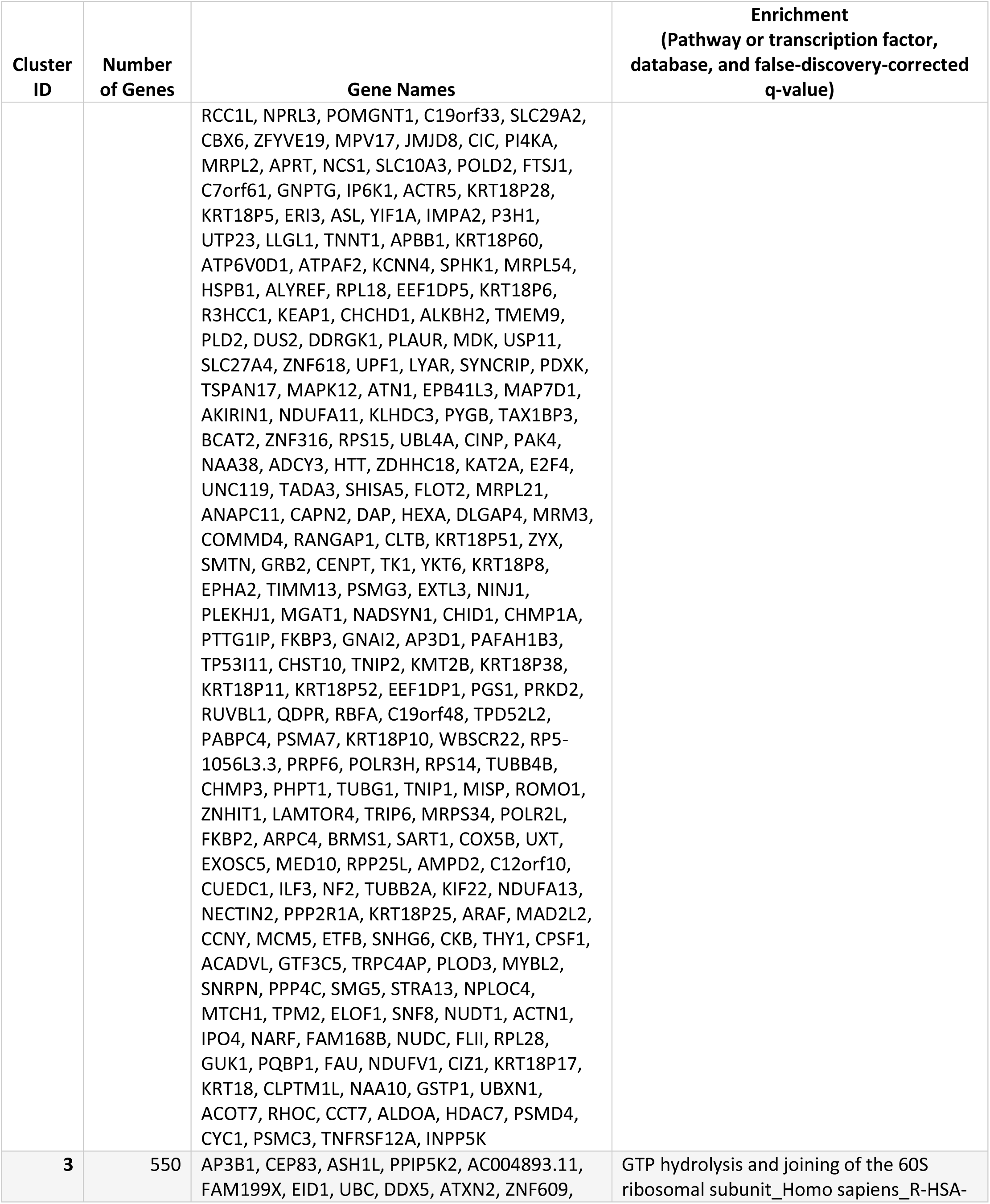
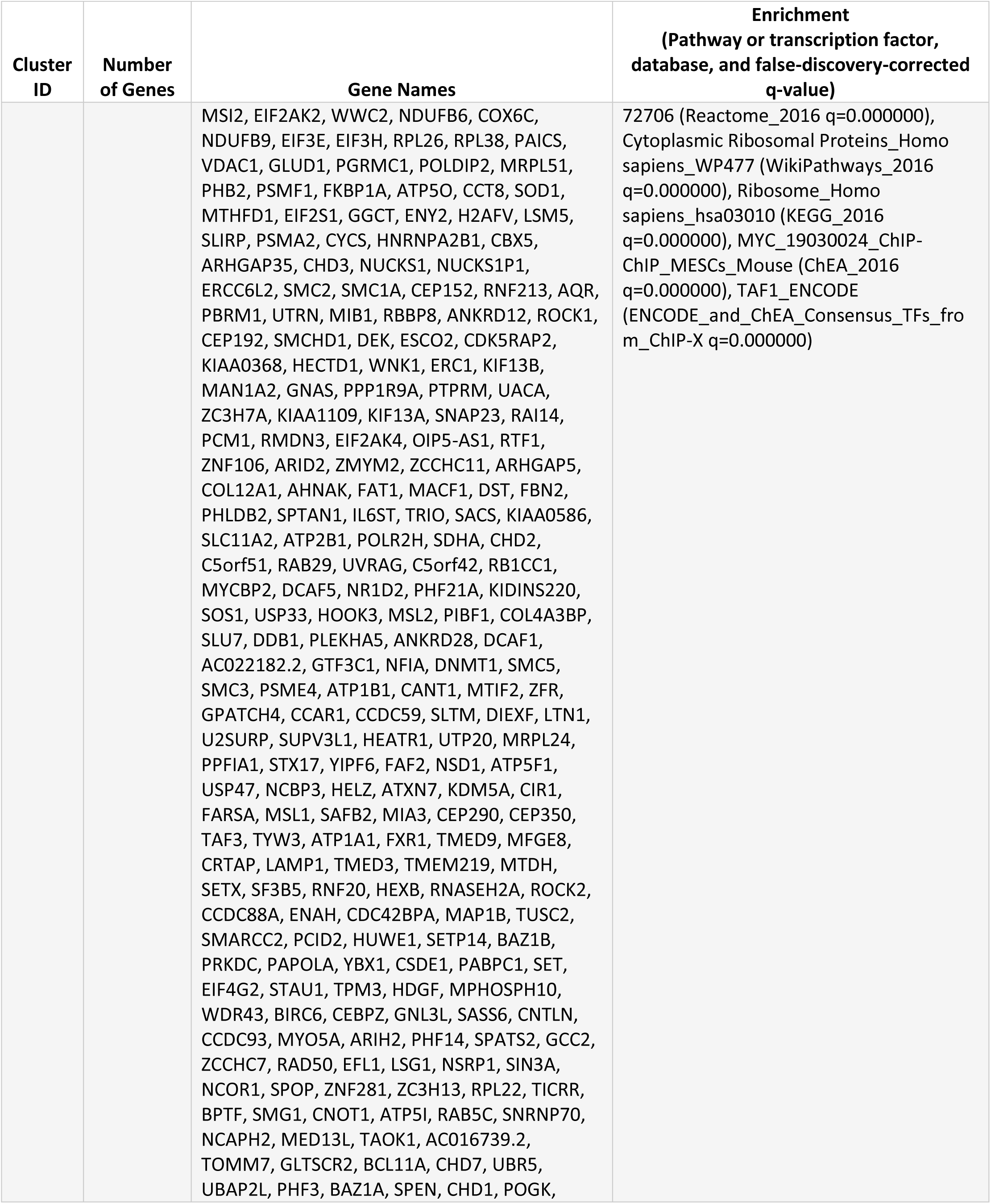
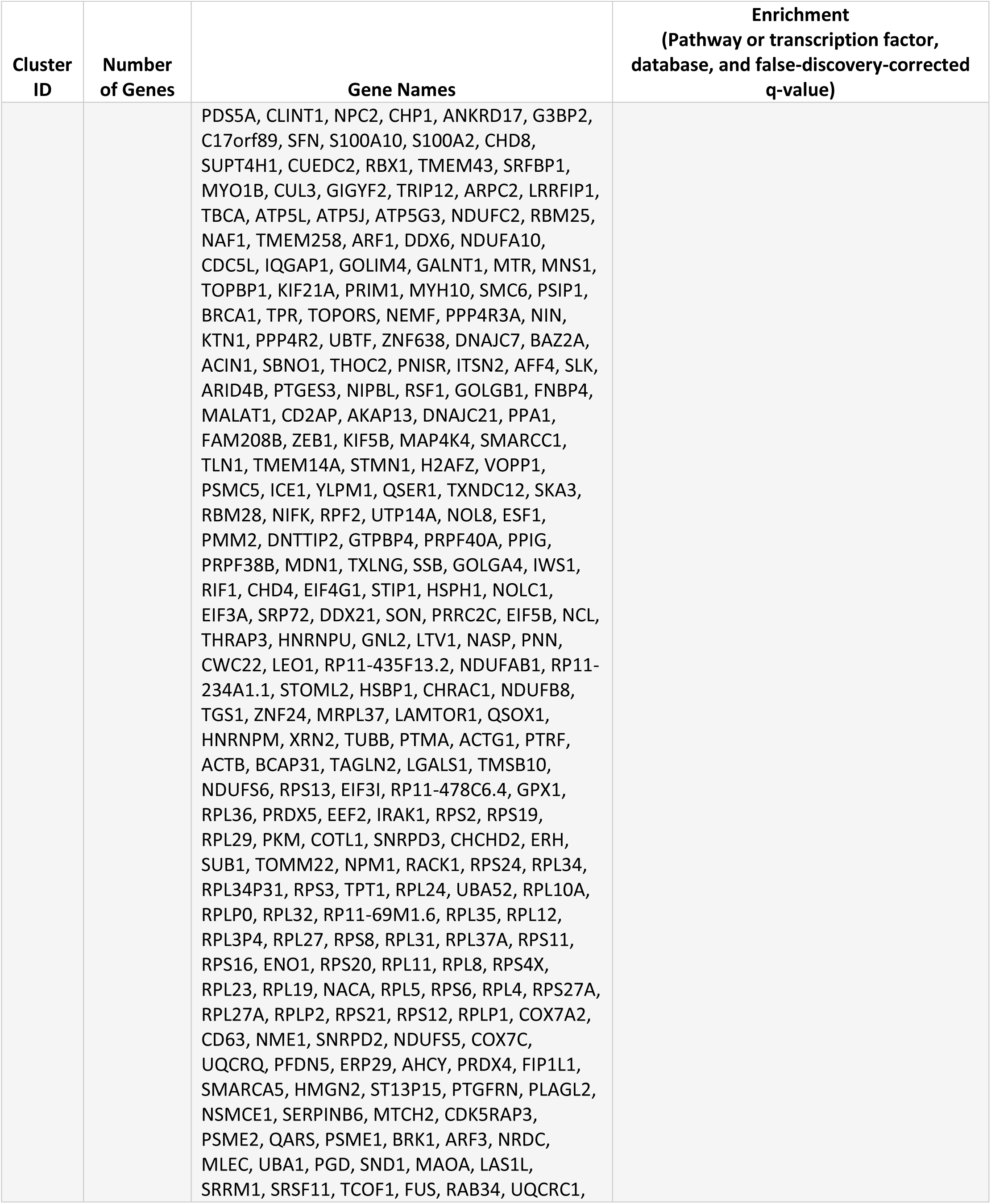
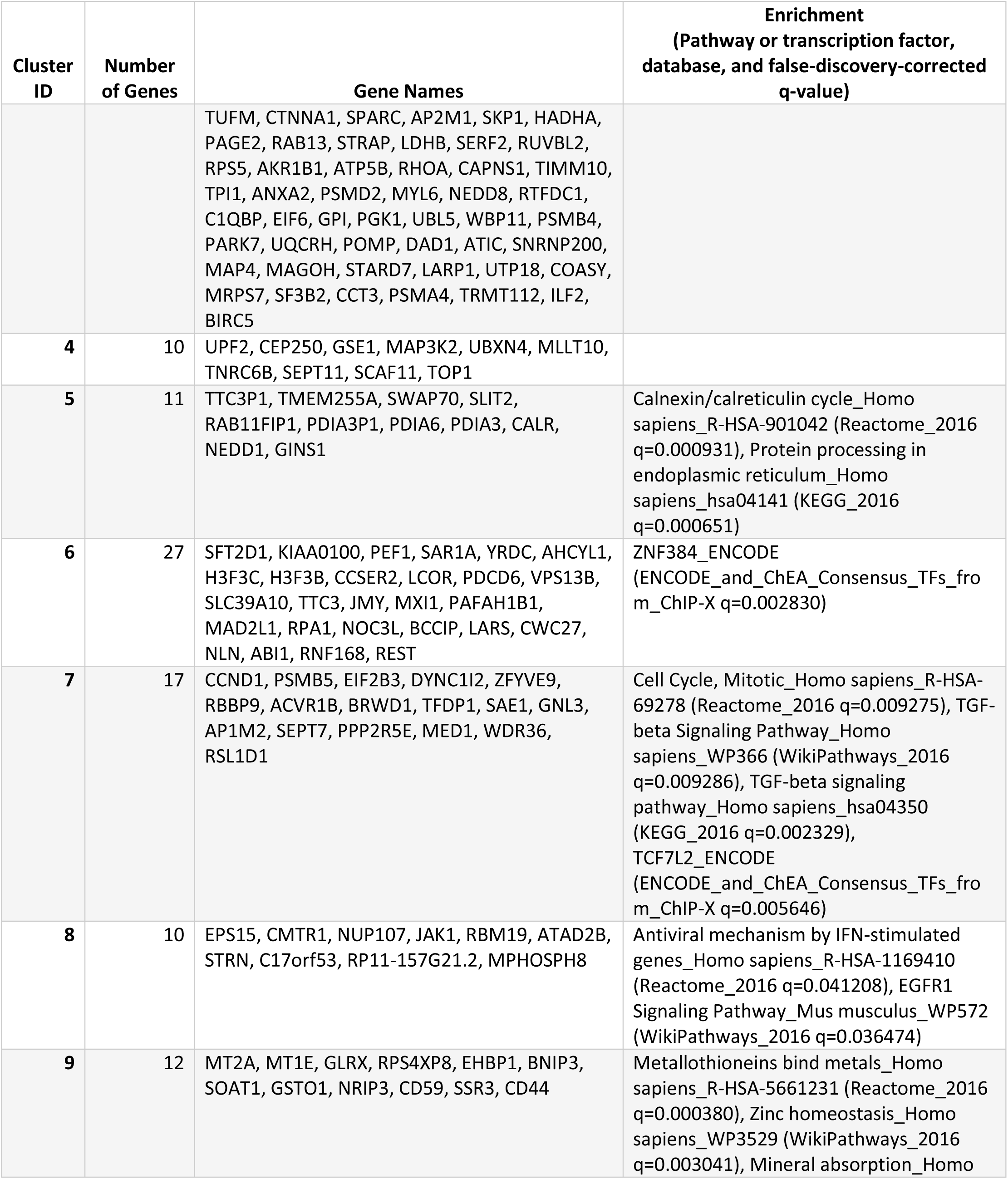
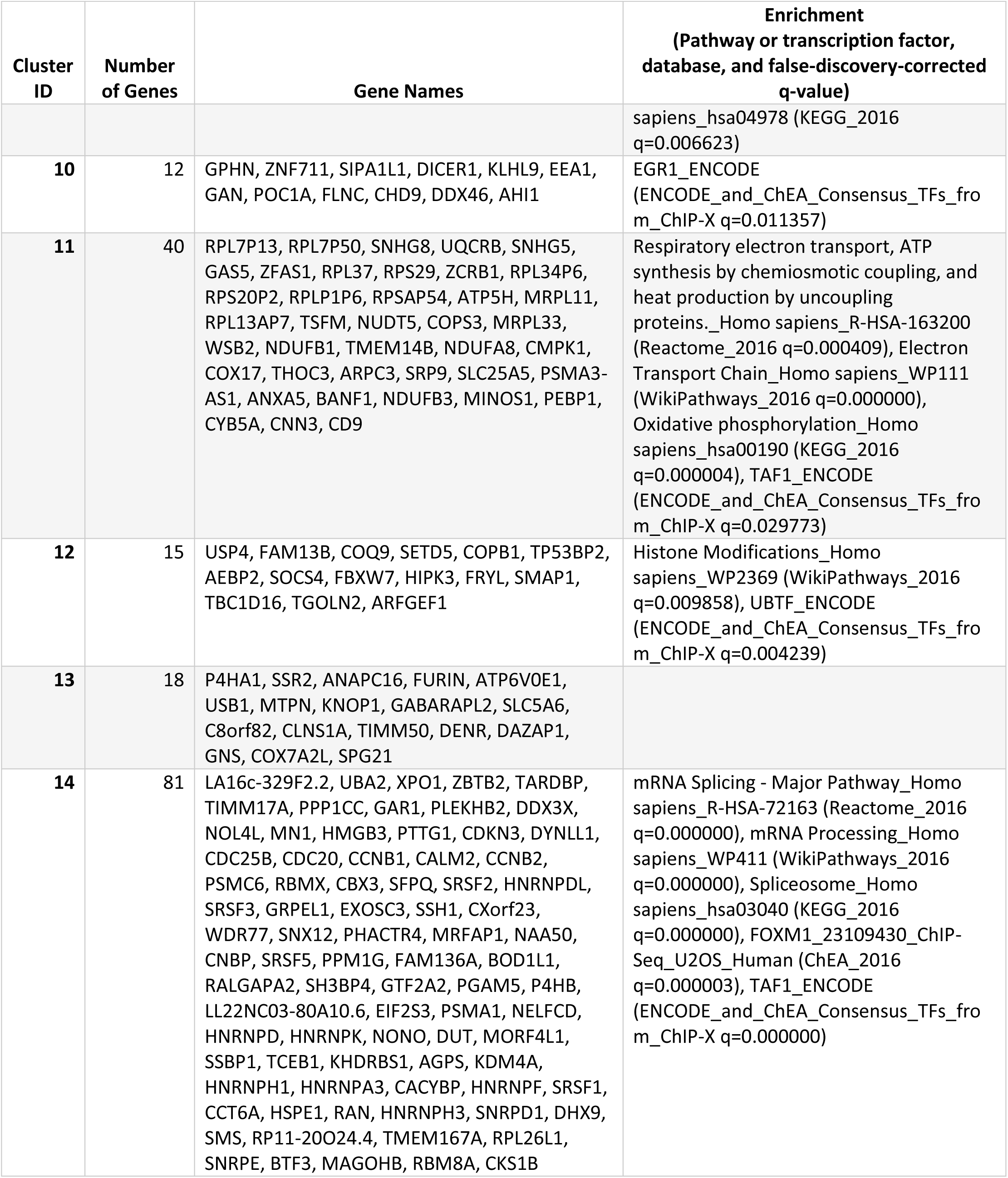
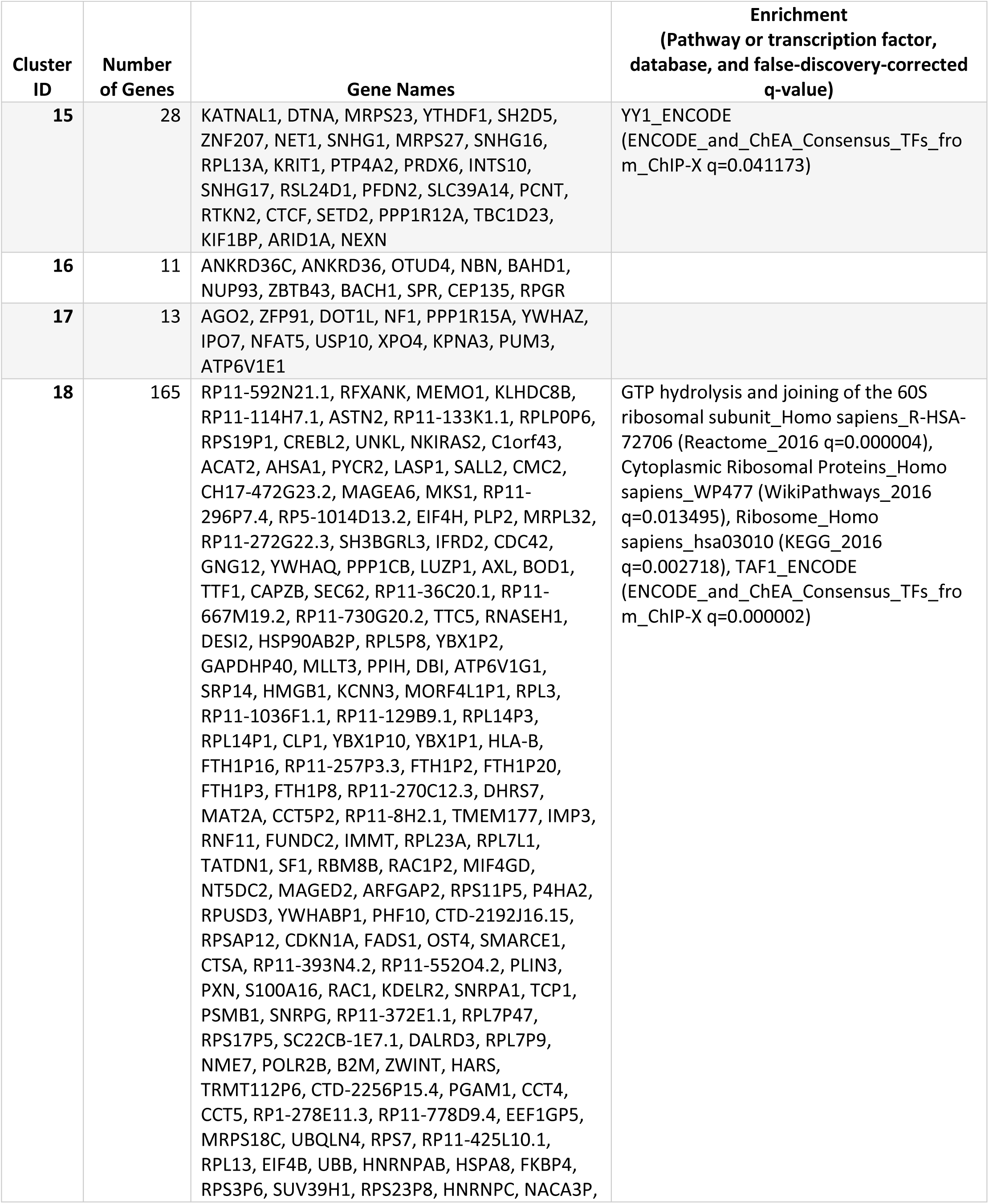
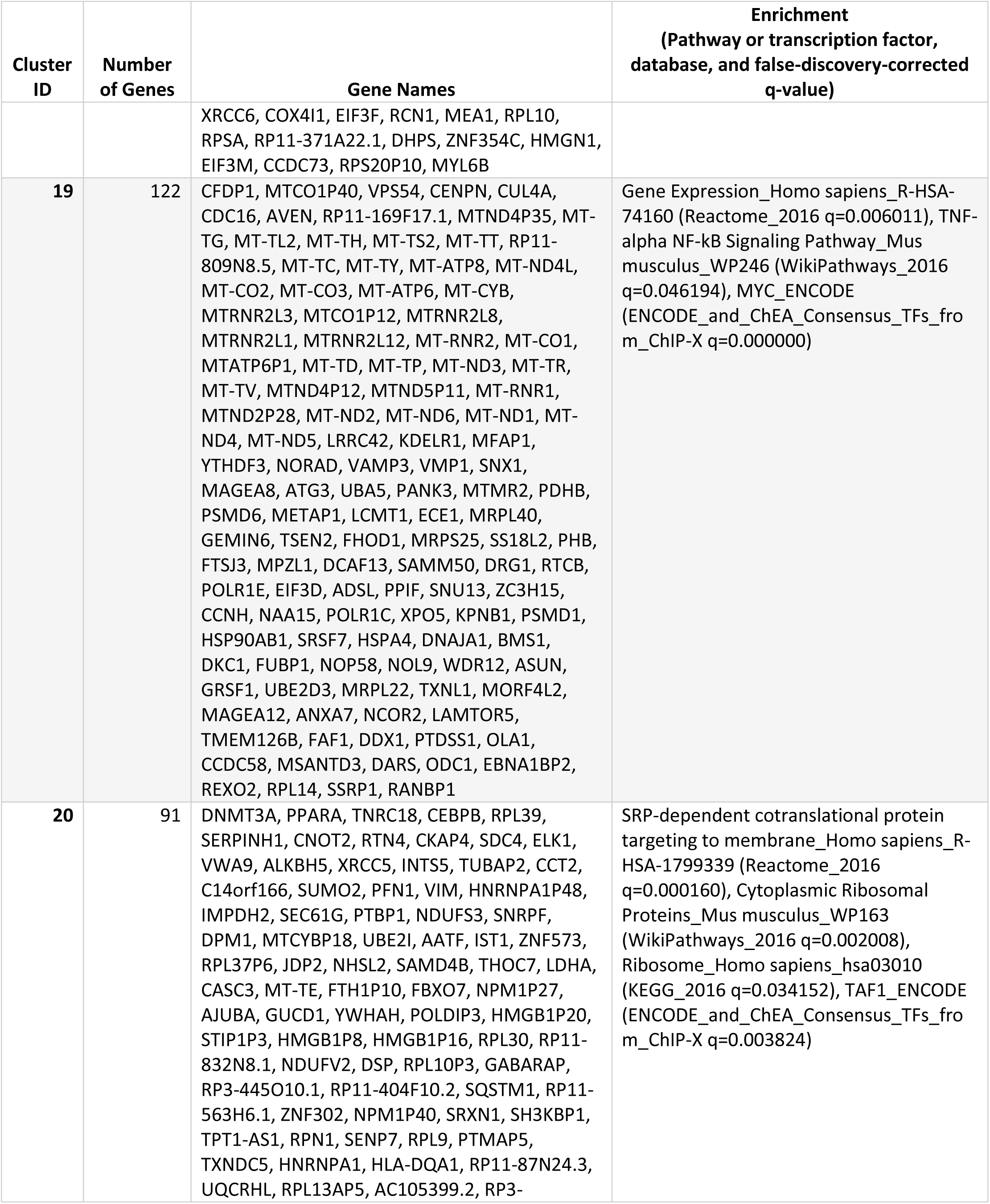
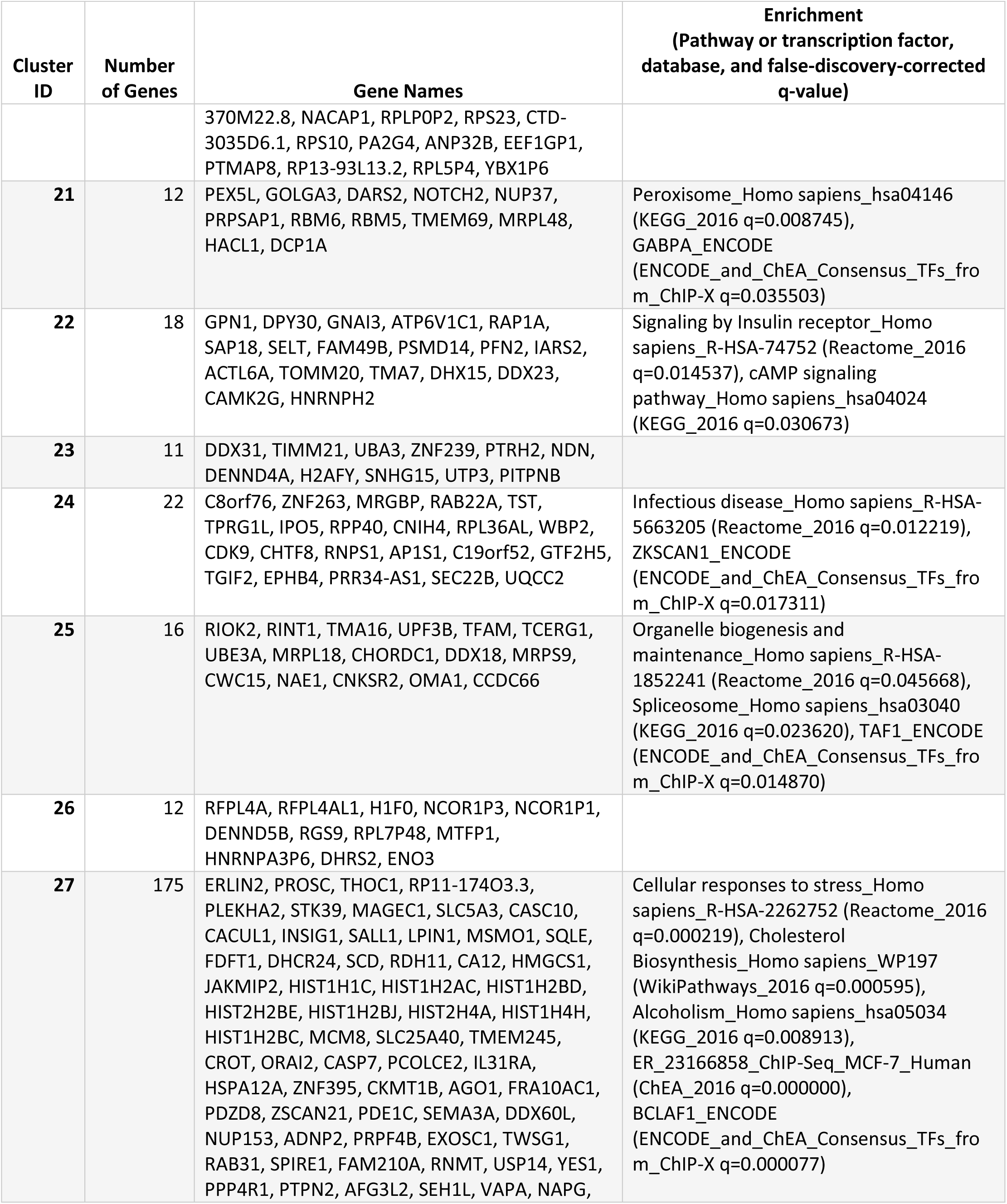
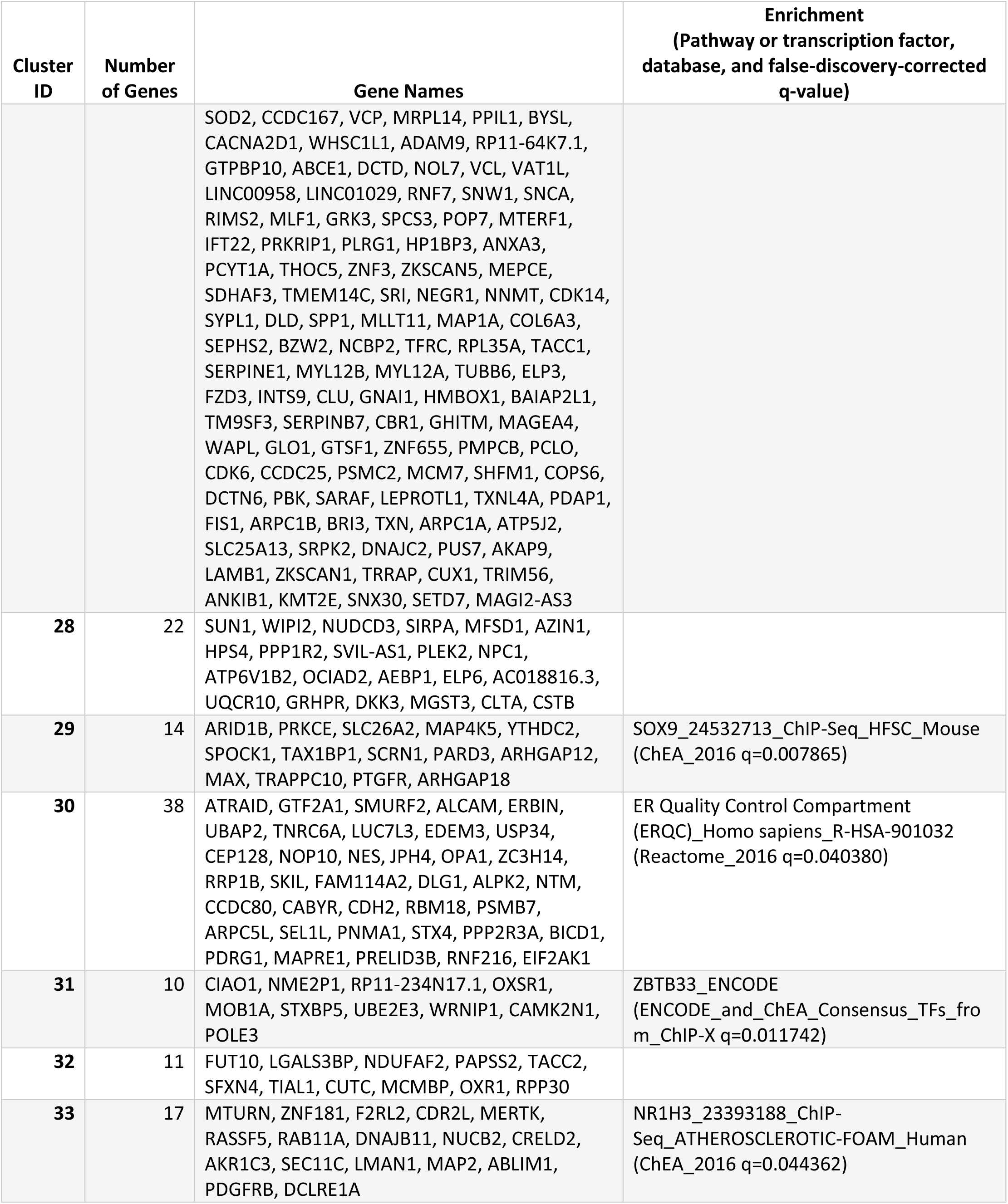
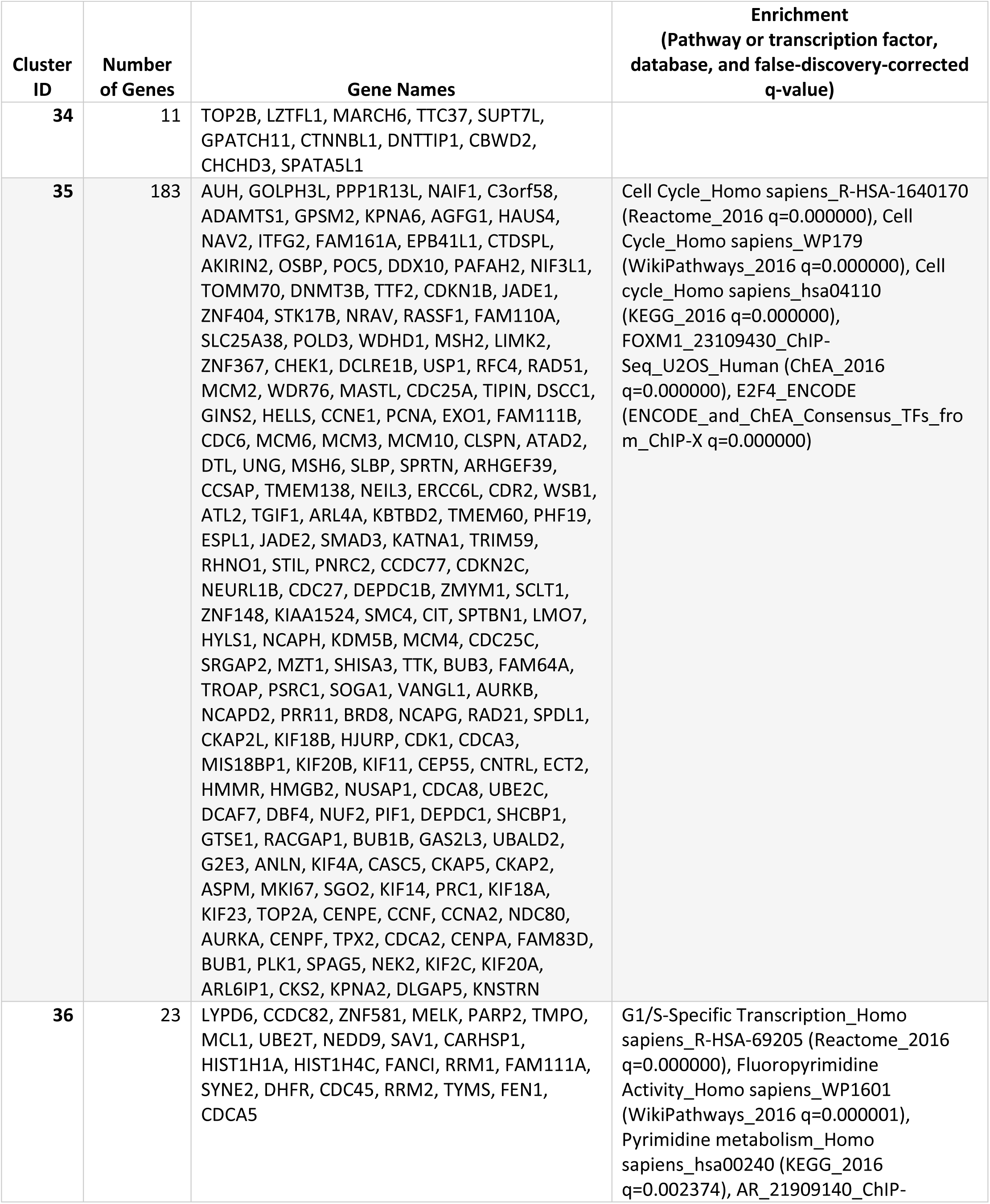
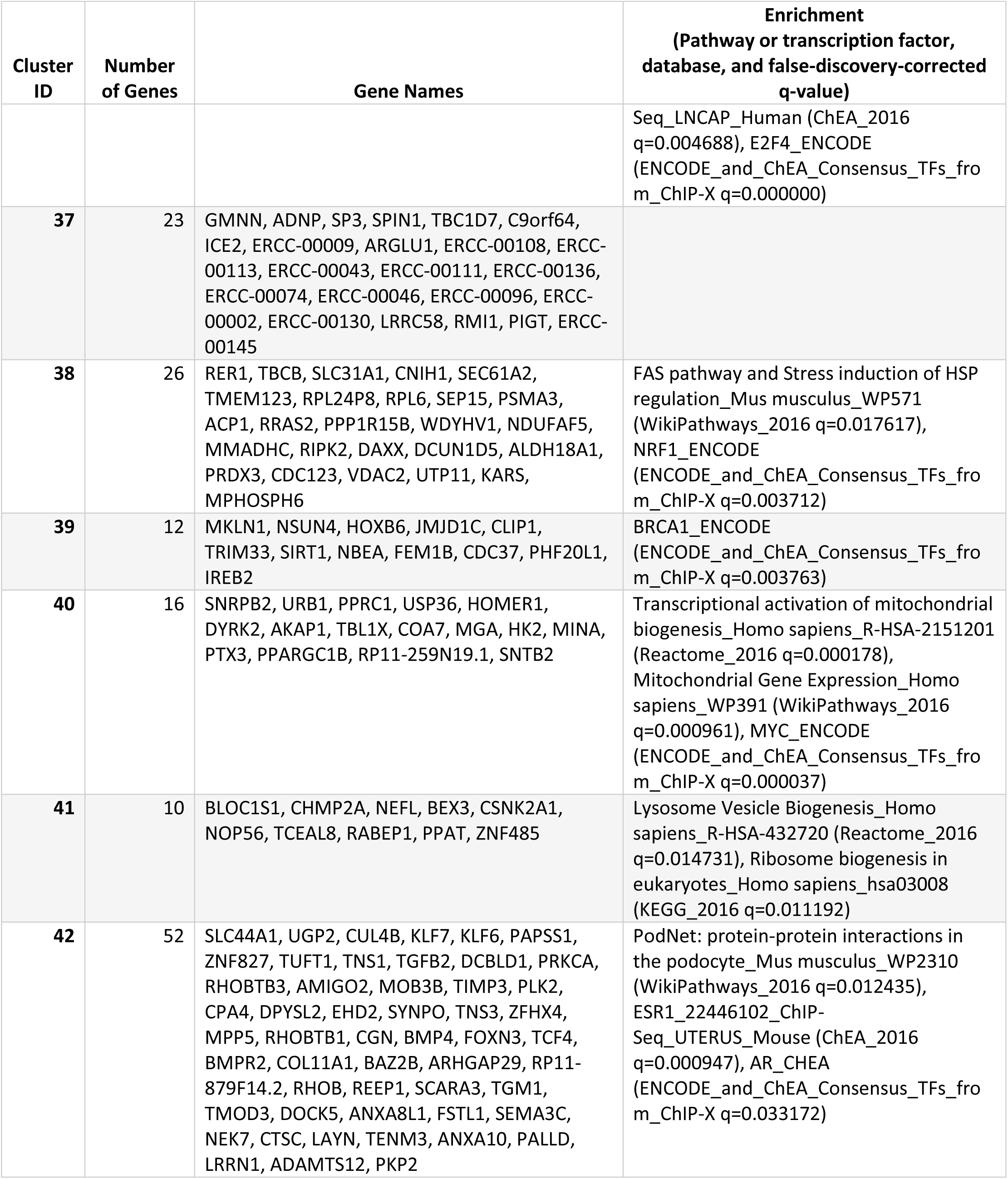
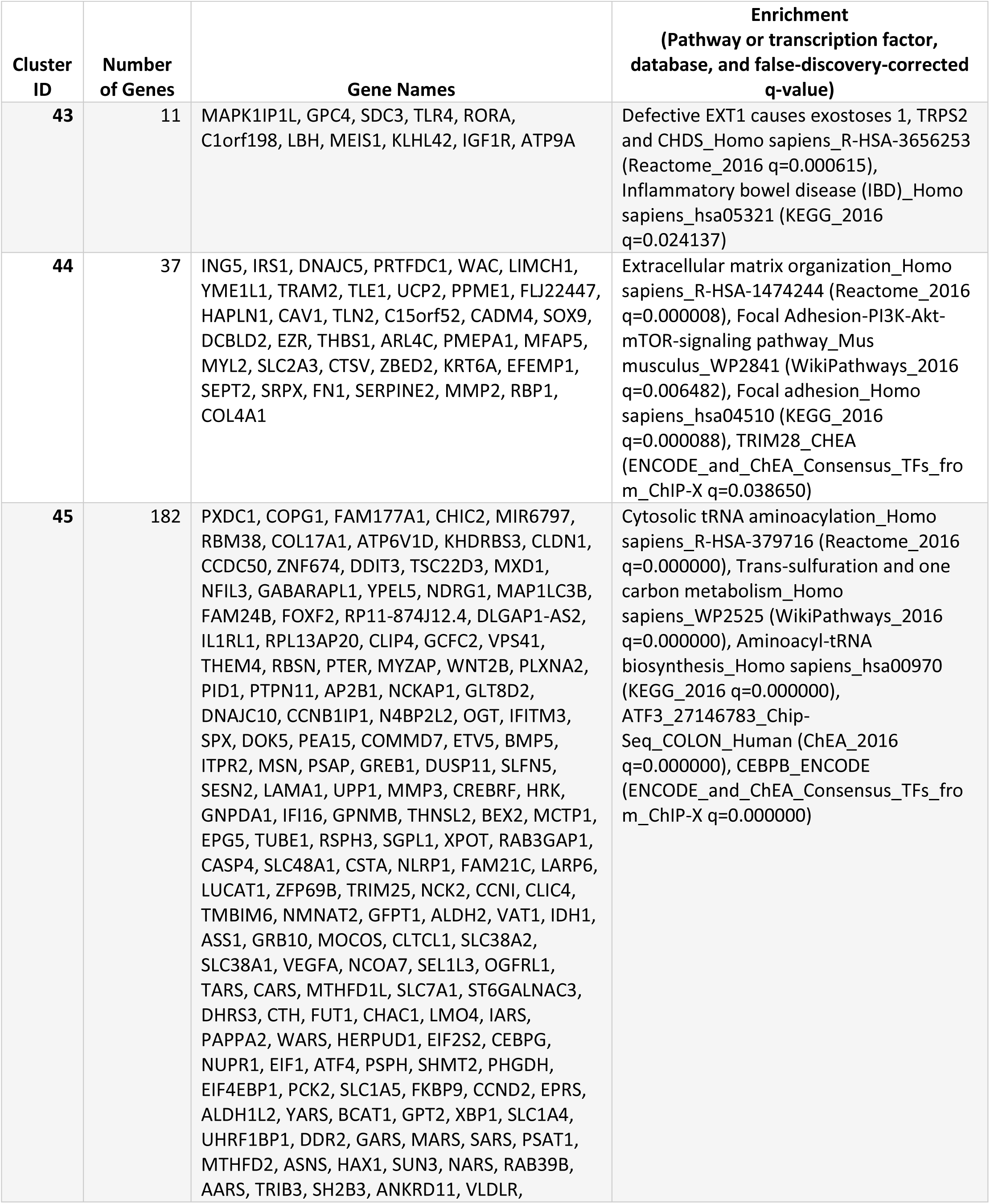
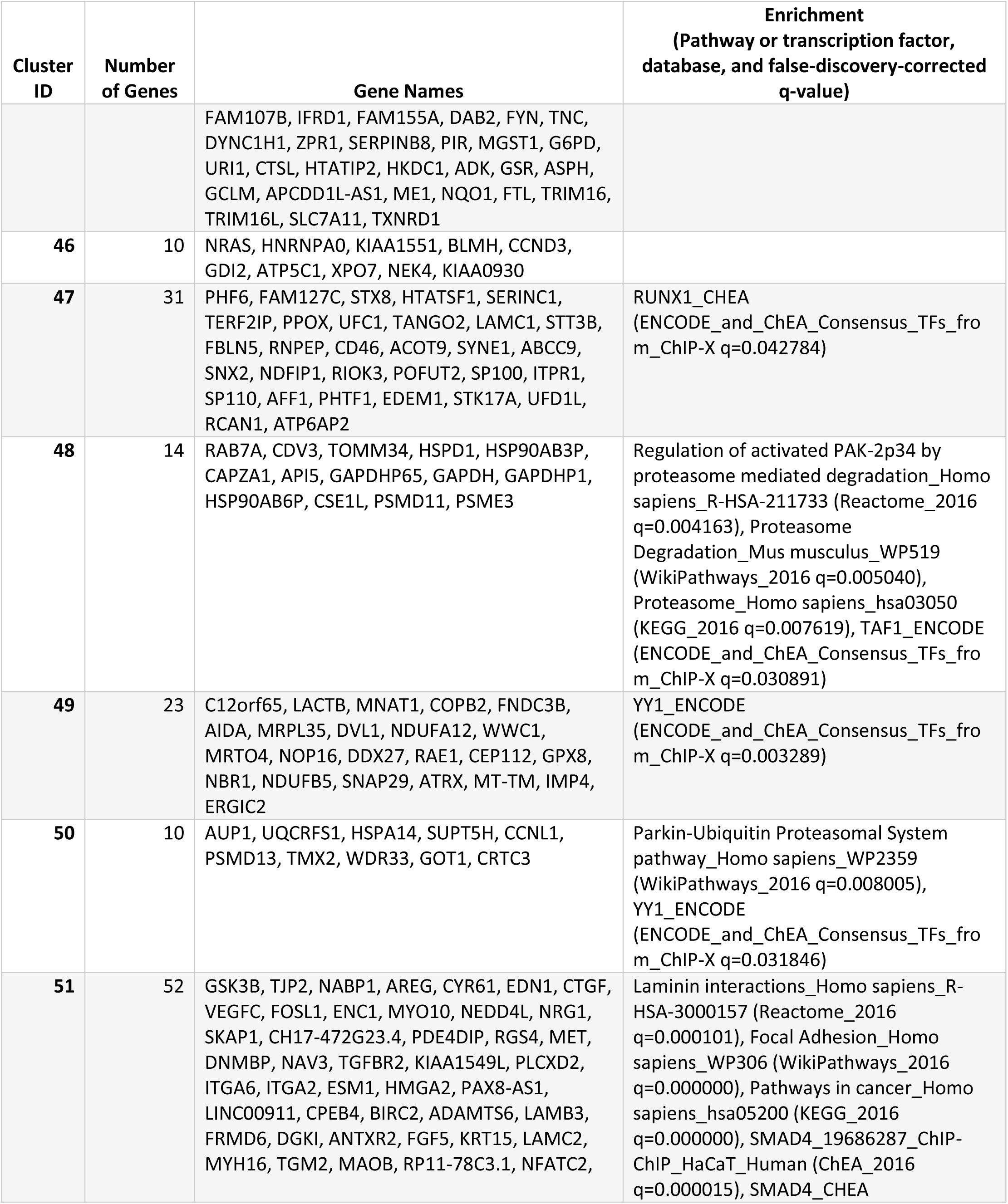
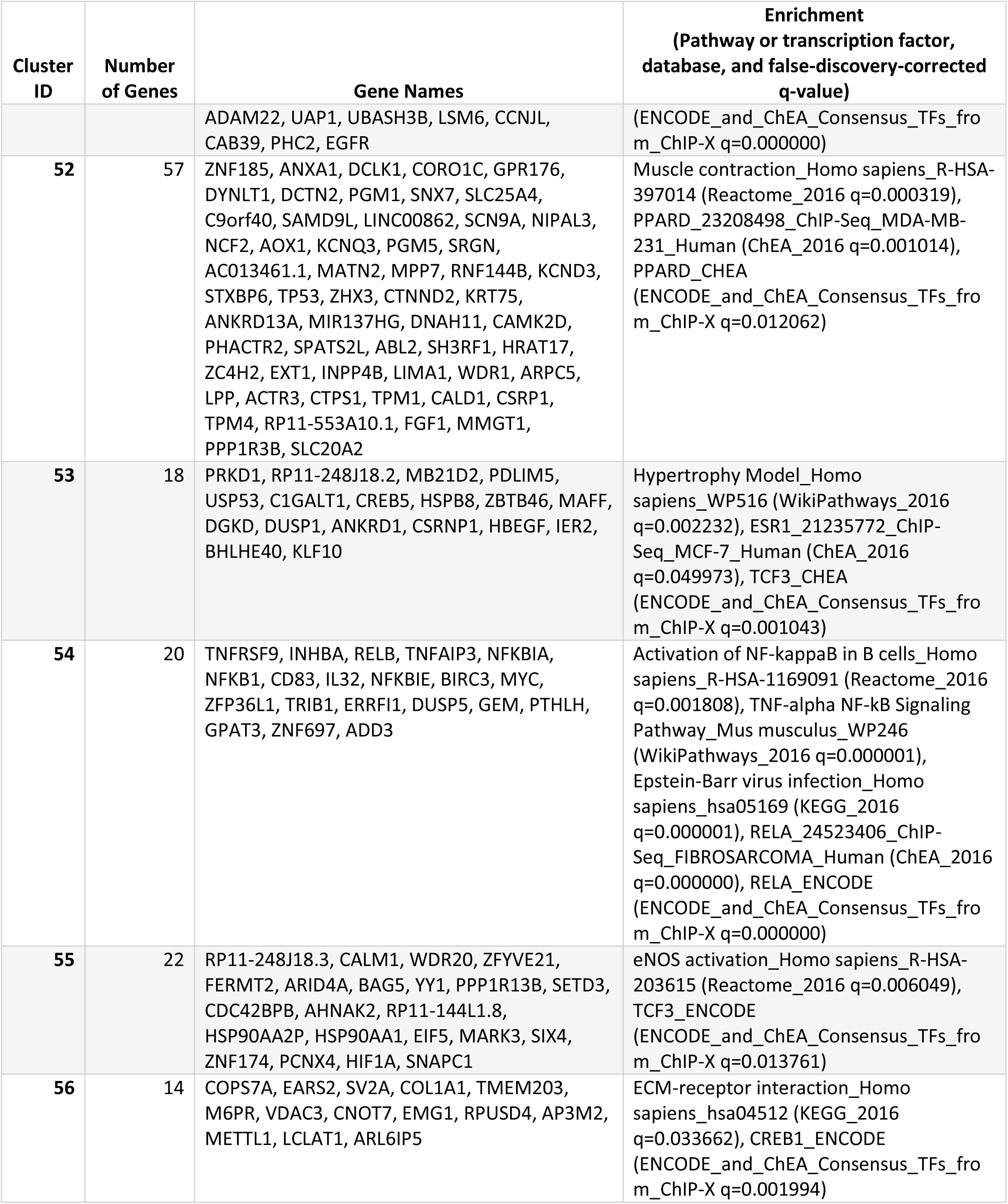
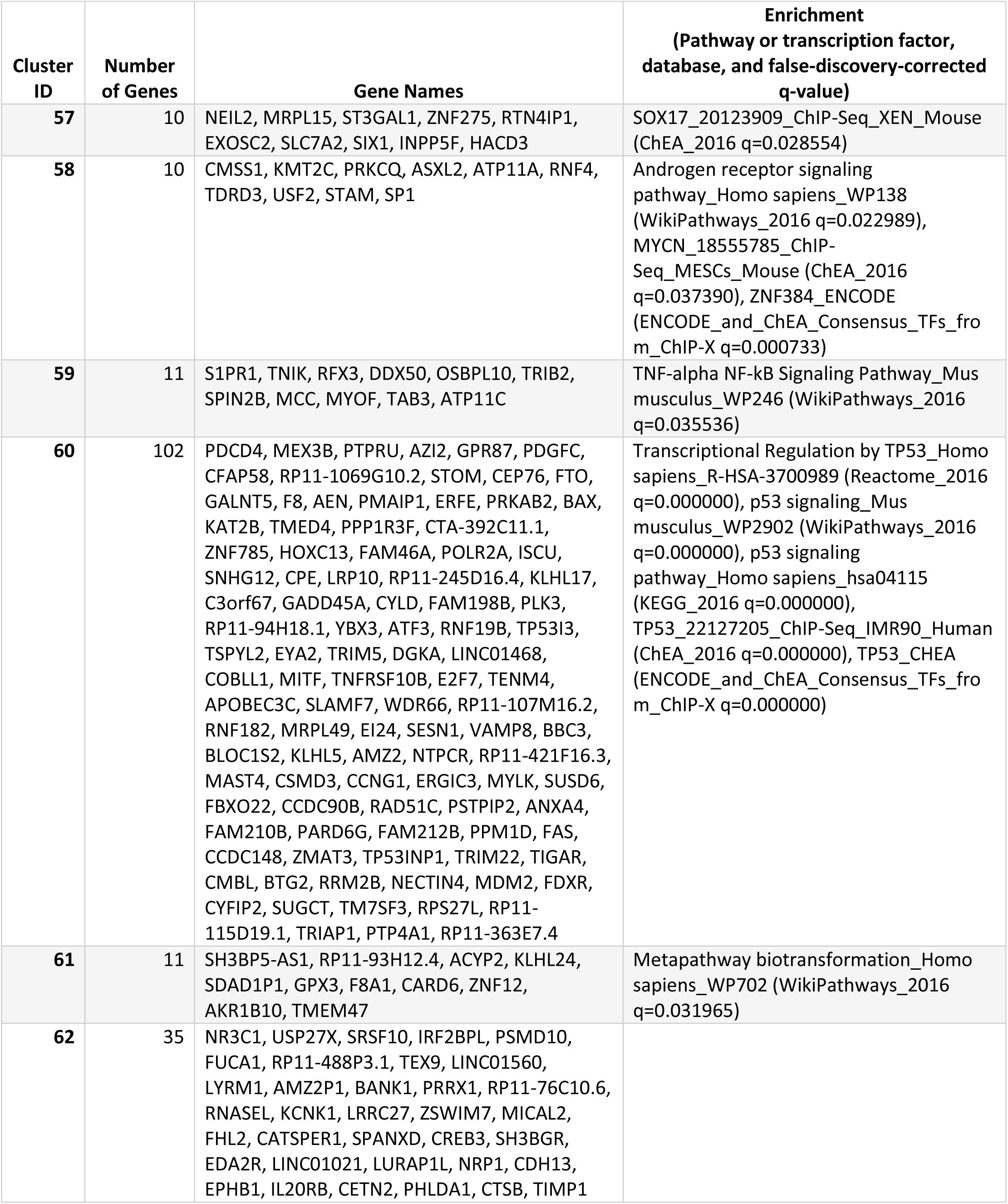
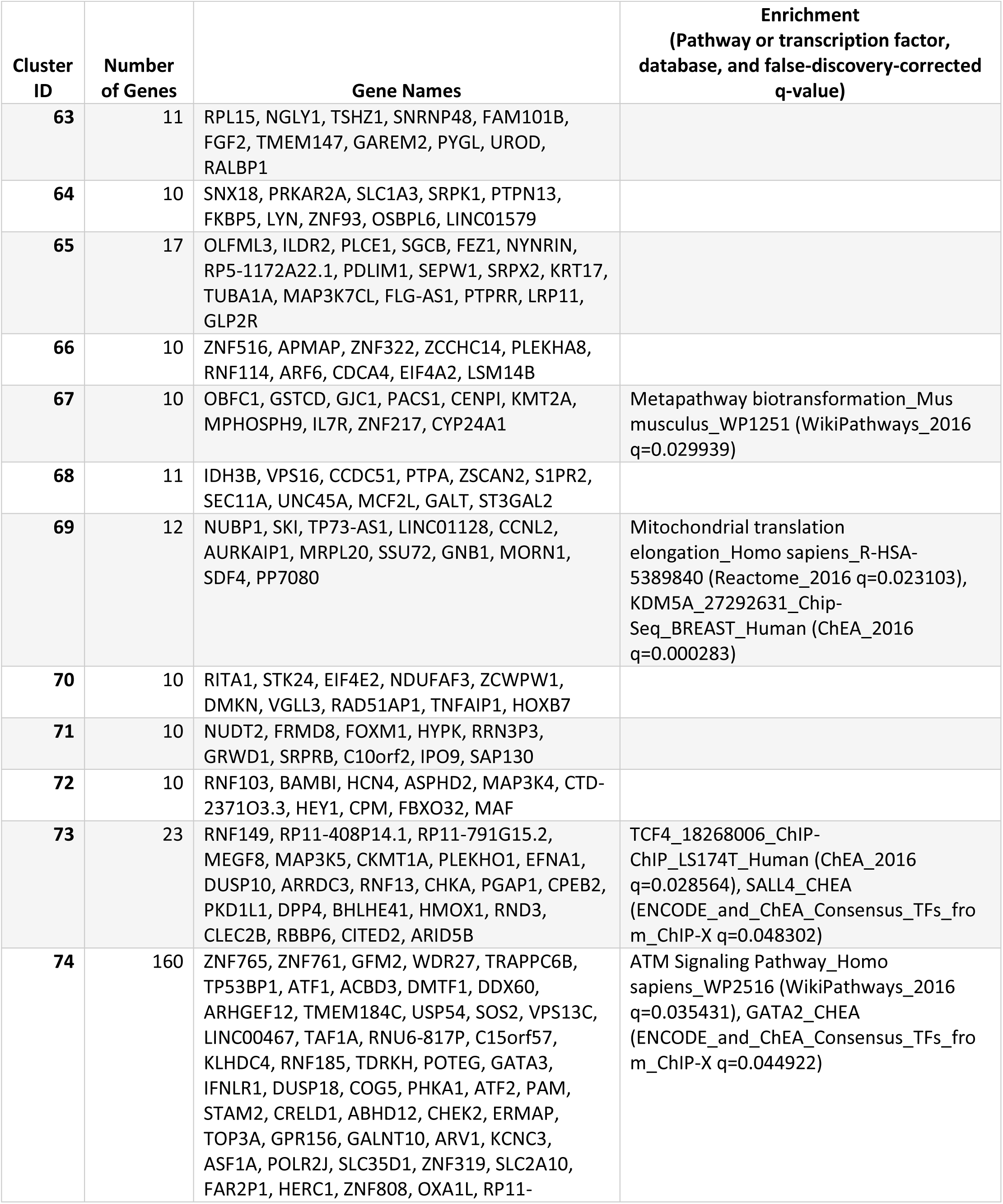
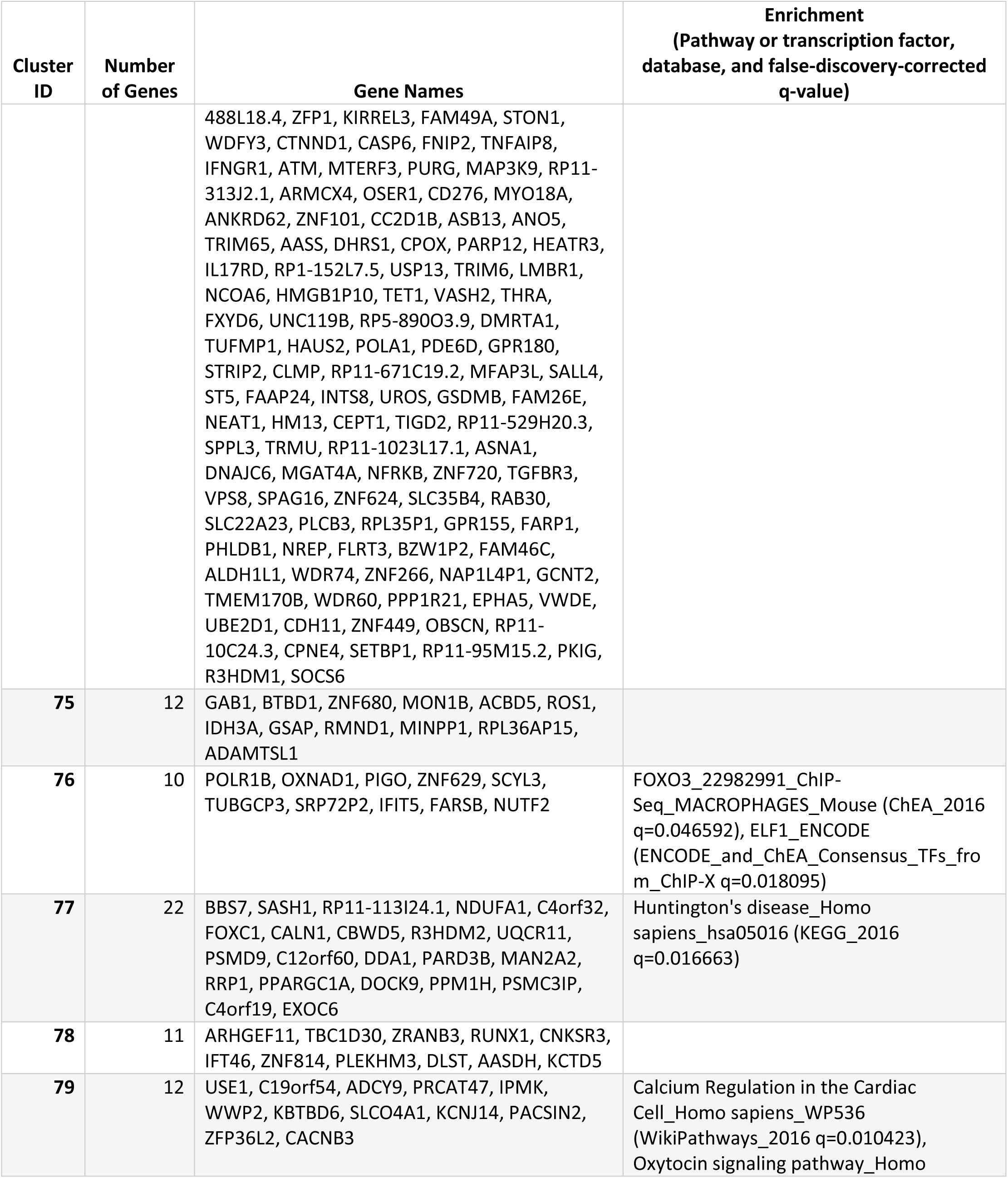
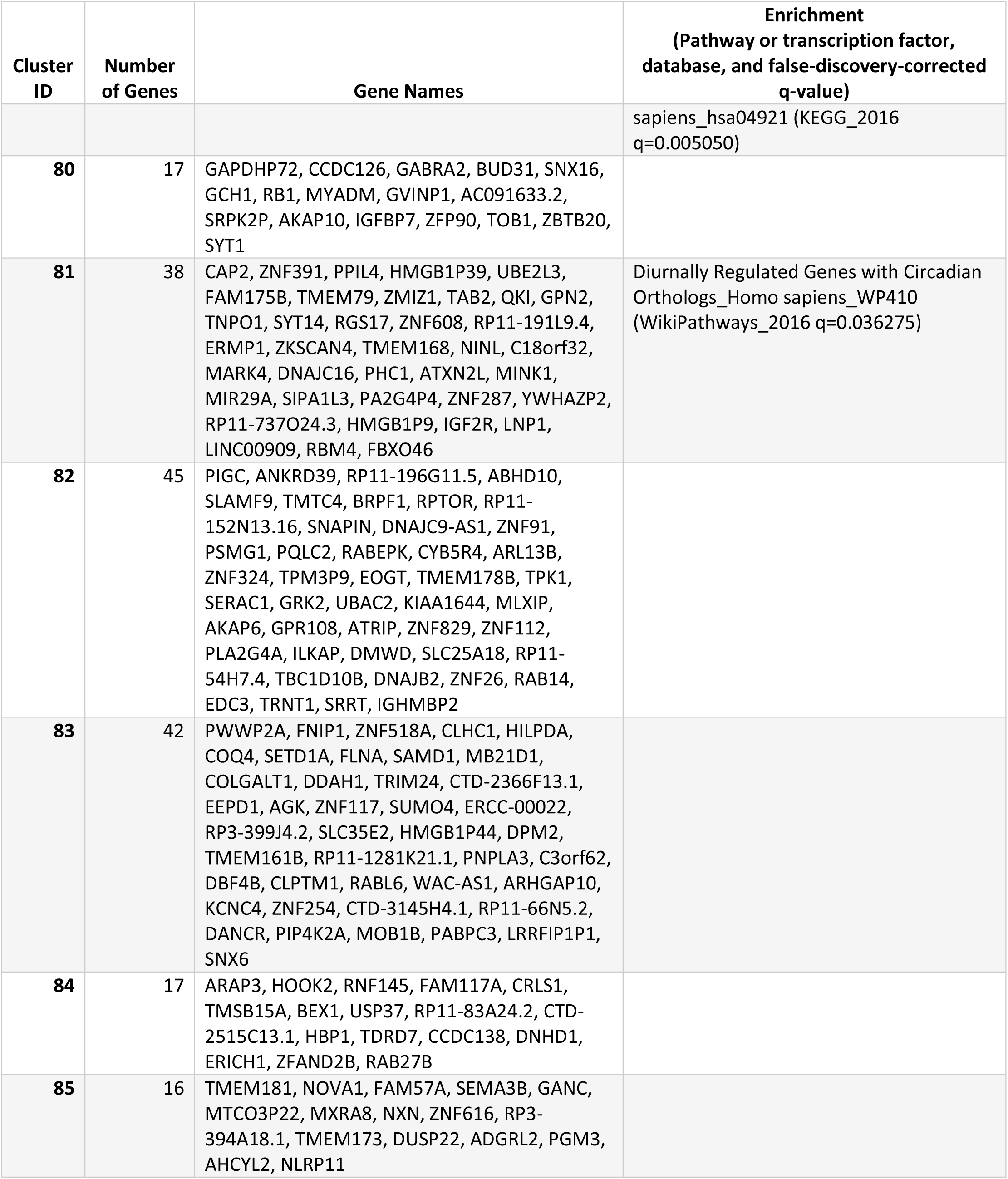
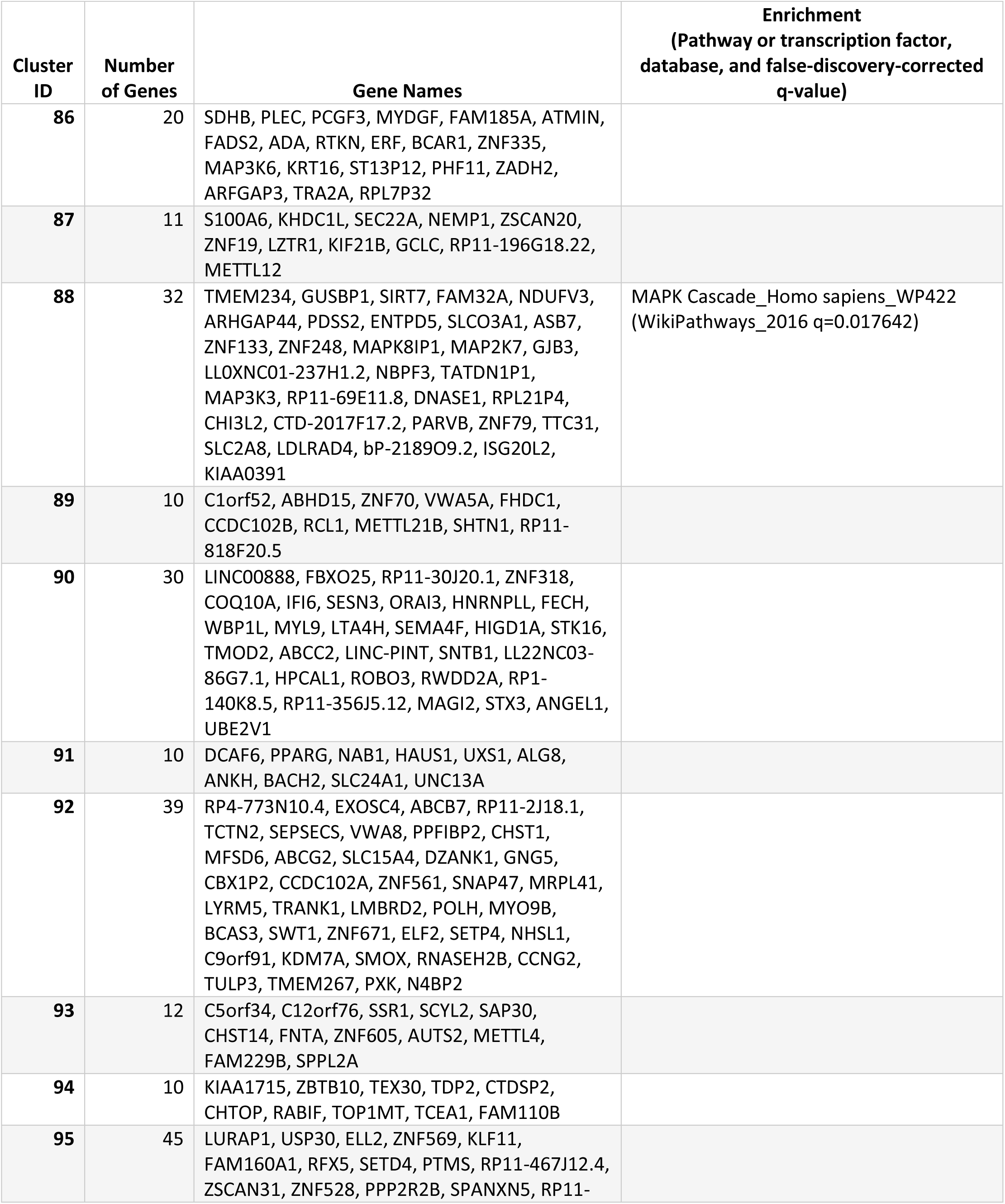
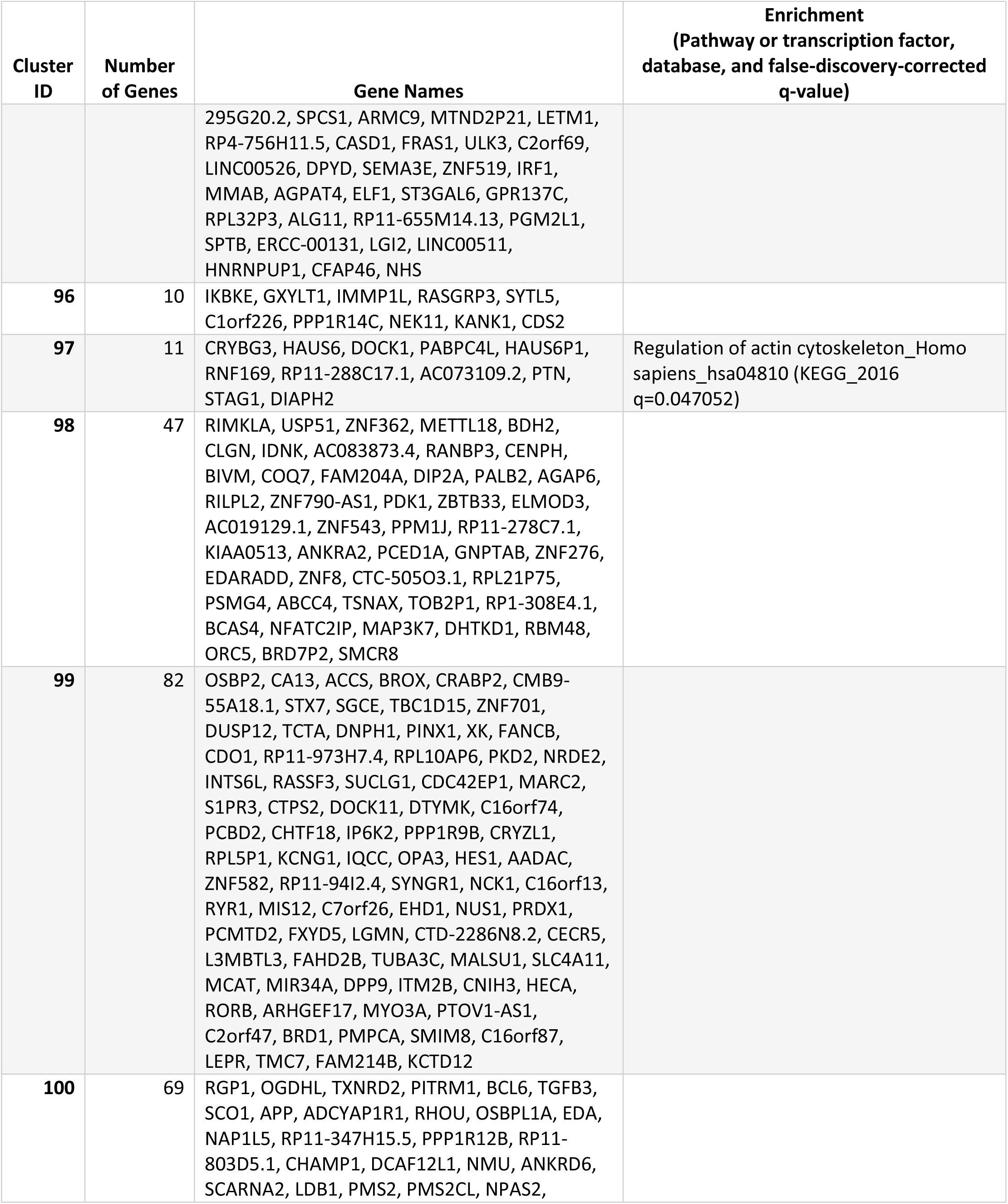
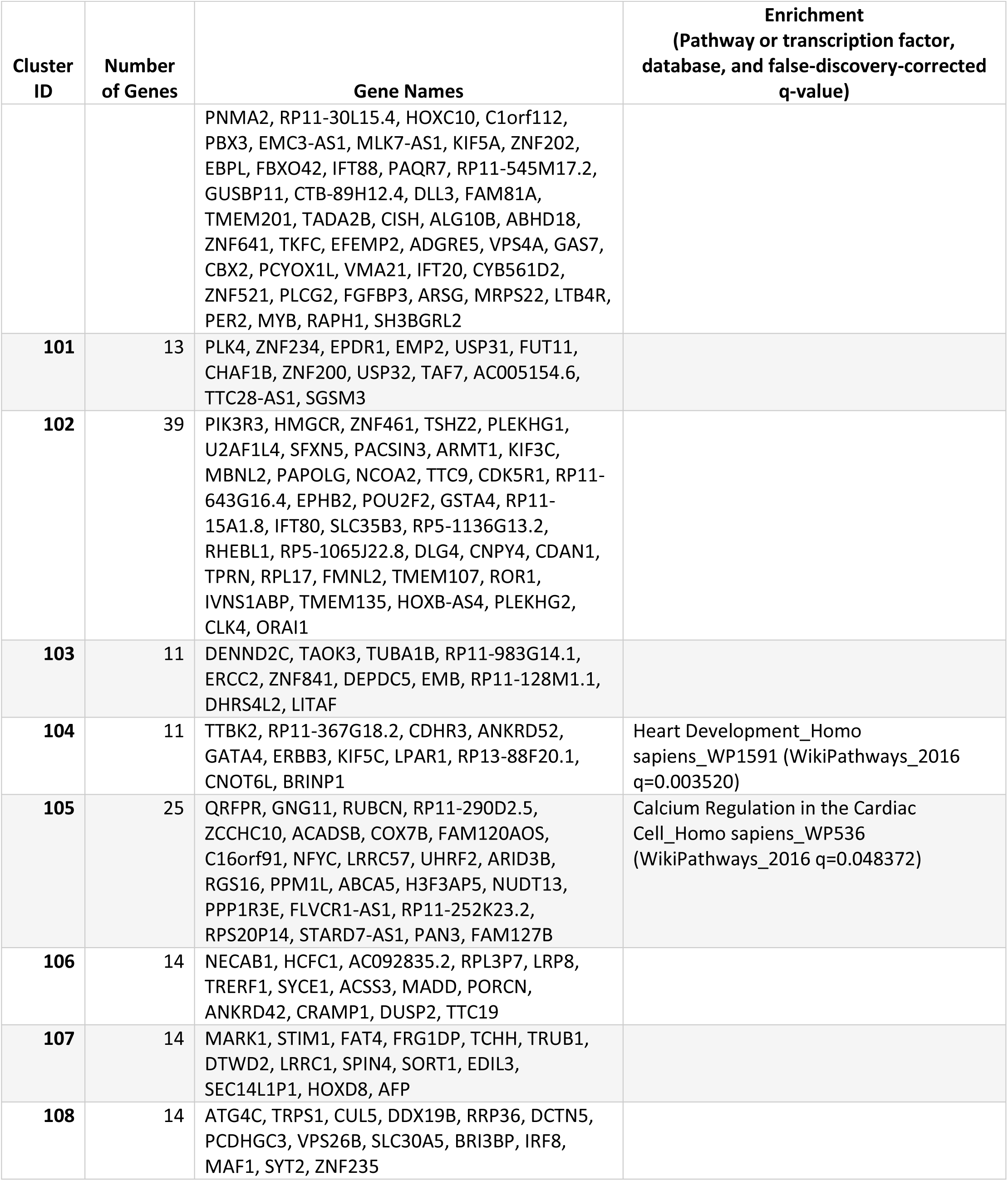
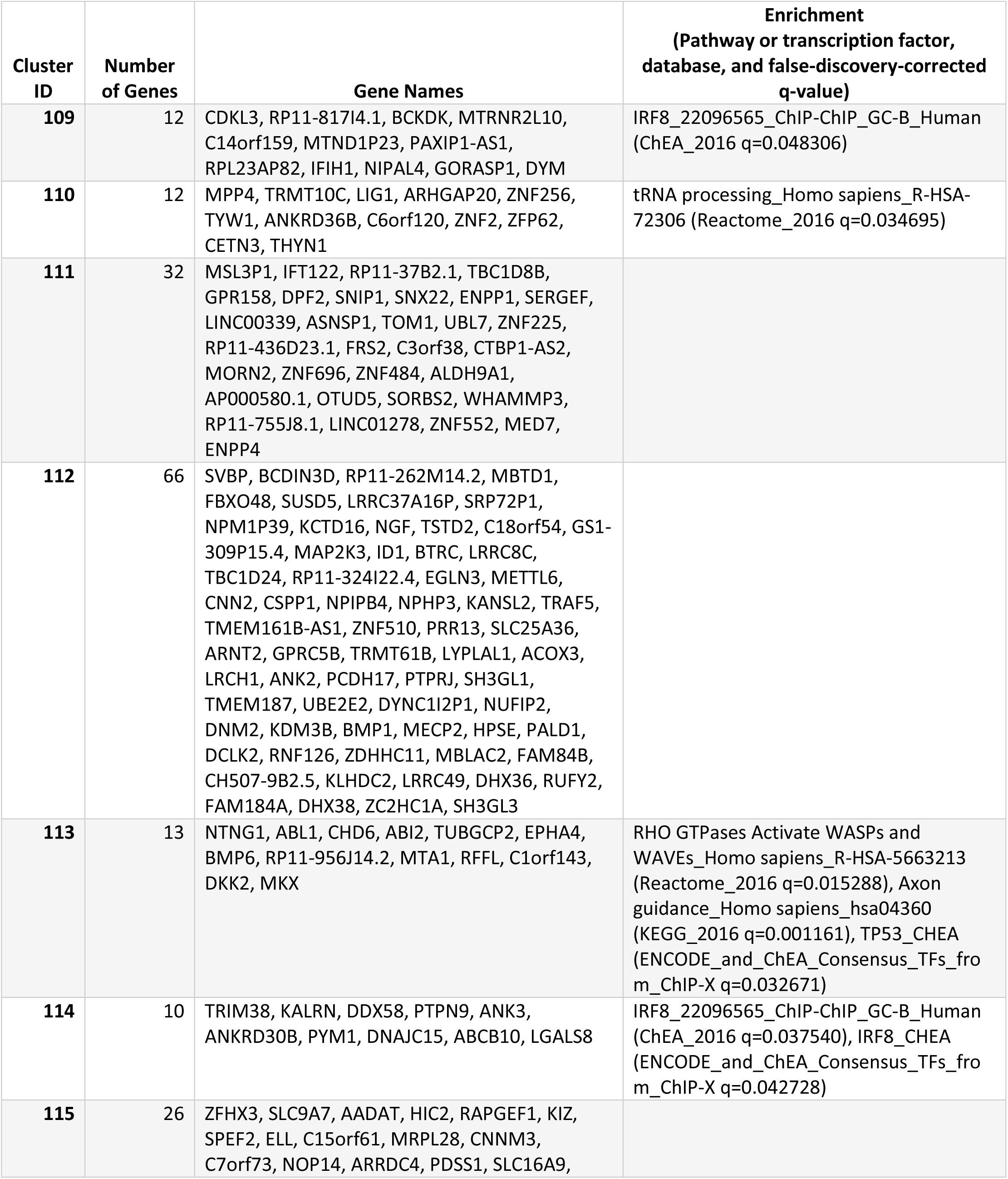
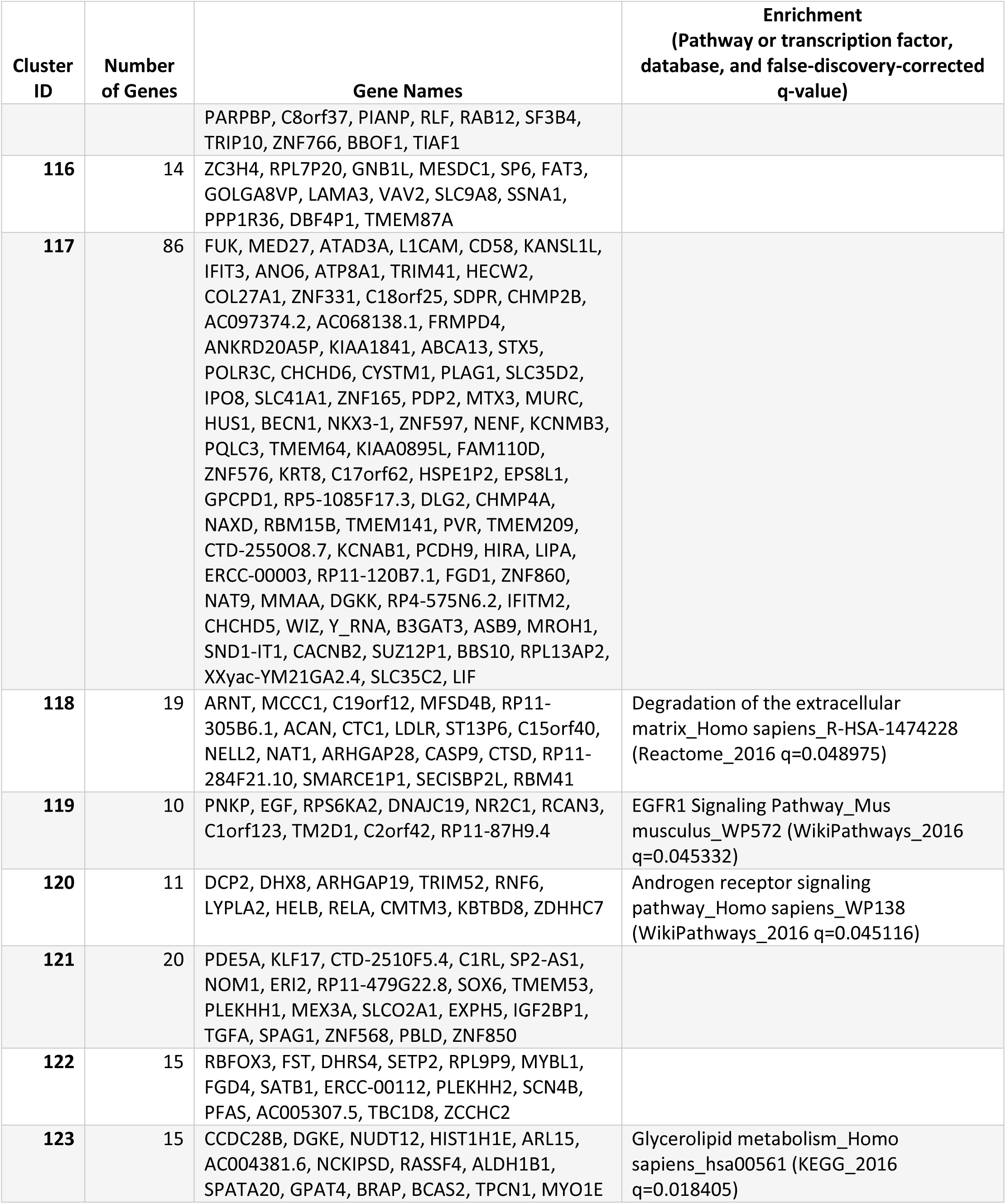
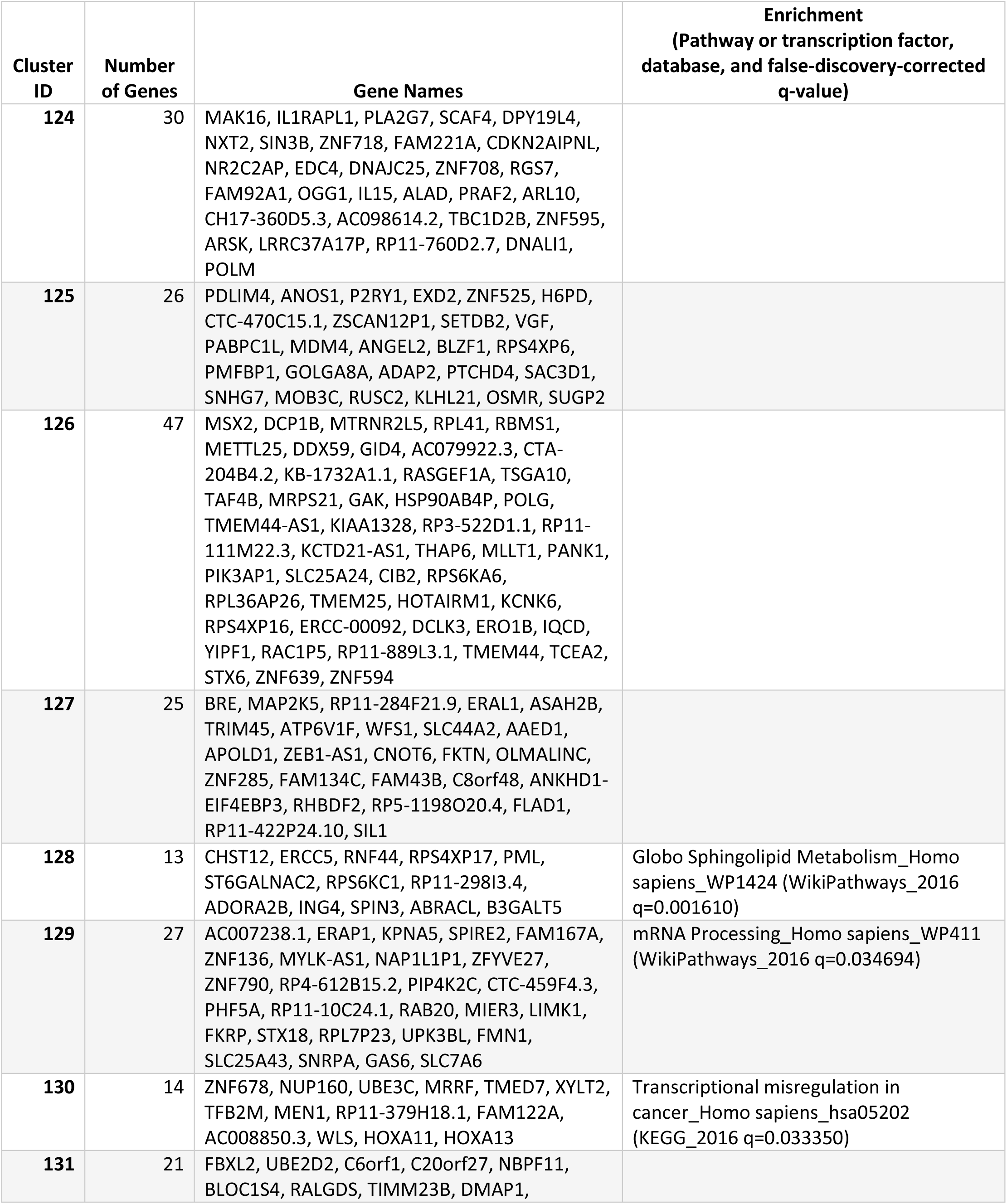
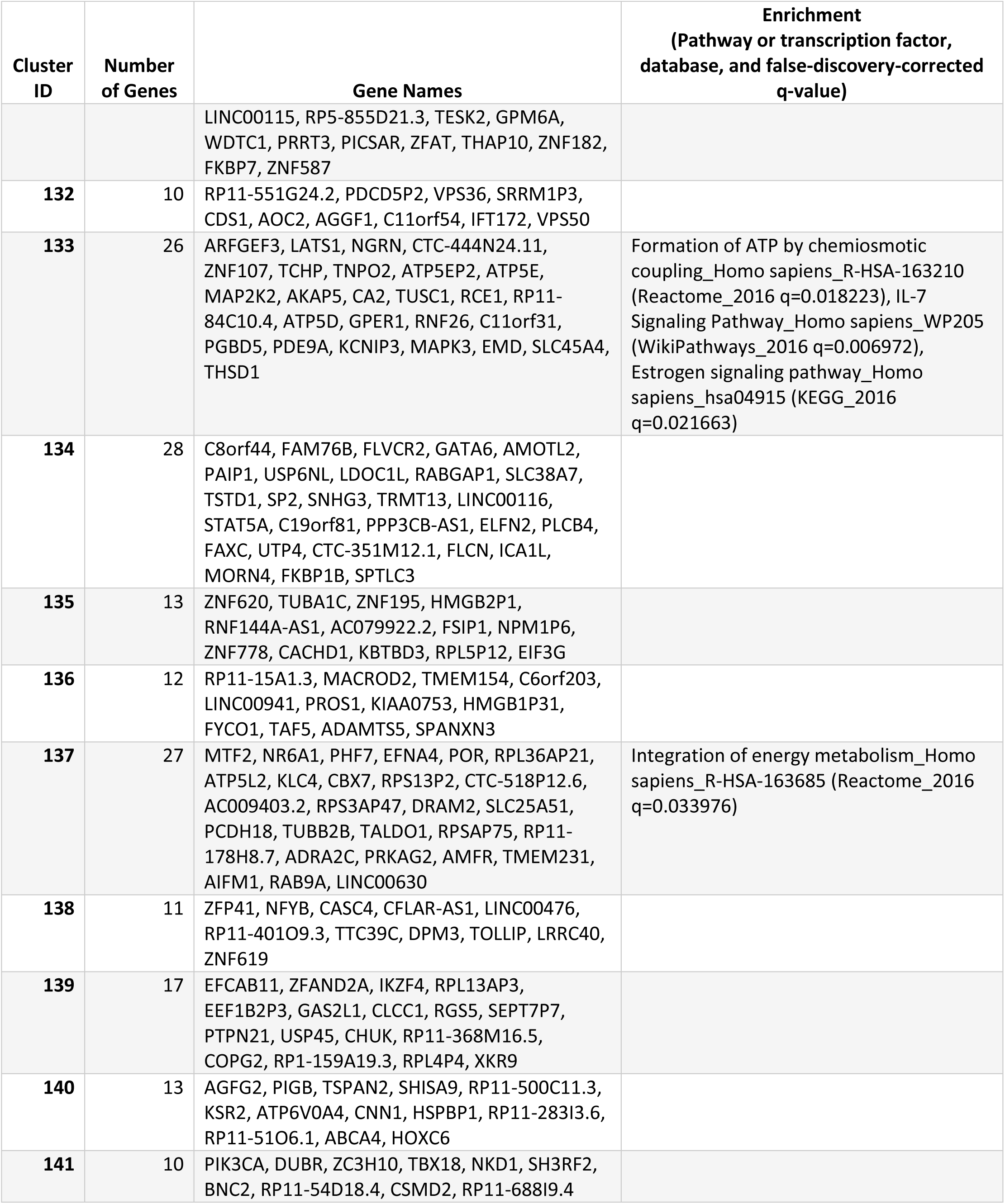
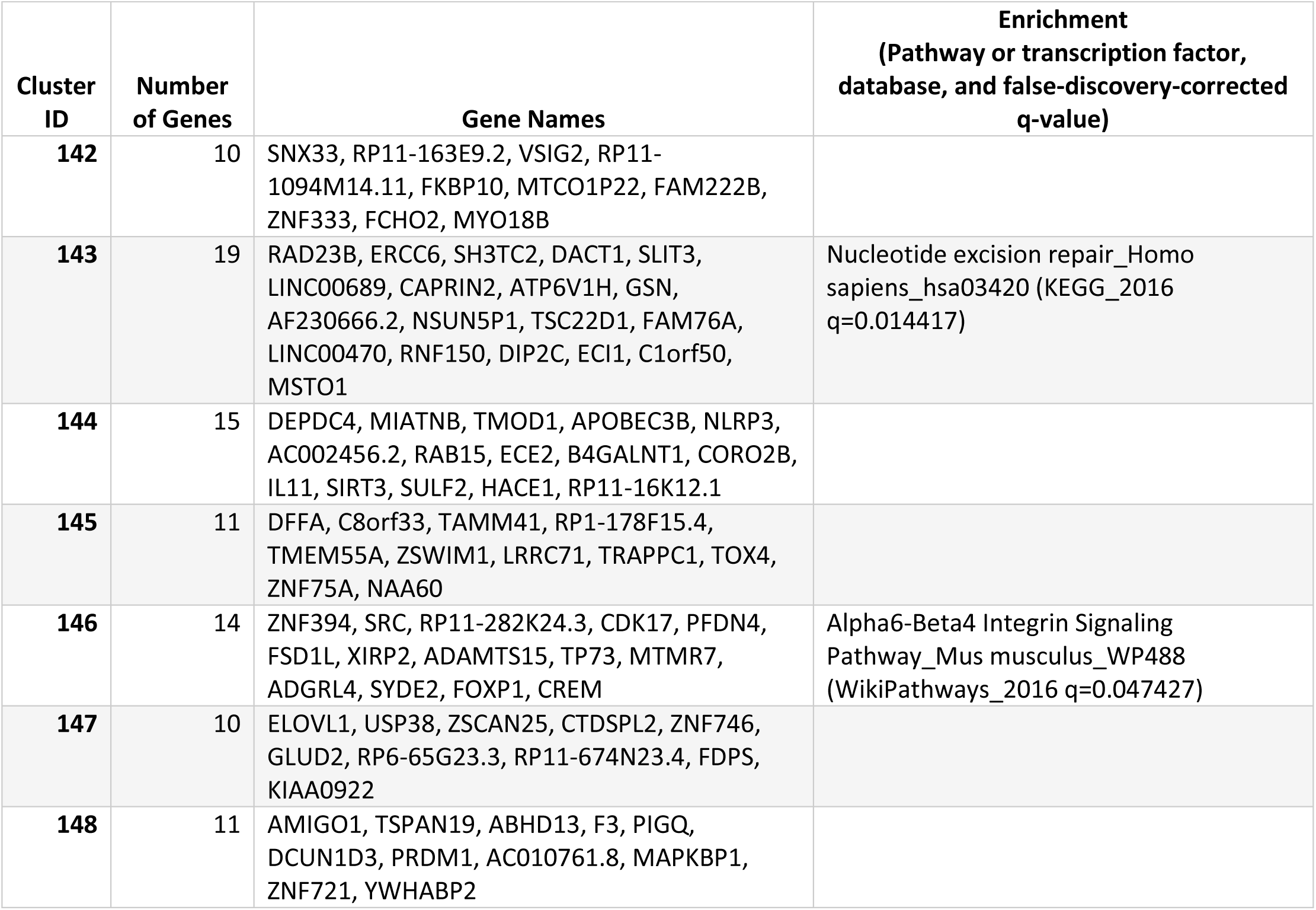
Functional enrichments of CGMs identified in U2OS. For each of the 148 CGMs identified in the U2OS dataset, the set of genes comprising the CGM is presented along with the enriched pathways and transcription factors identified by Enrichr (*35, 36*).

